# Finding a Balance: Characterizing Teaching and Research Anxieties in Biology Graduate Teaching Assistants (GTAs)

**DOI:** 10.1101/2021.09.10.459805

**Authors:** Miranda M. Chen Musgrove, Kate Petrie, Alyssa Cooley, Elisabeth E. Schussler

## Abstract

Graduate students in the United States are reporting increased anxiety, affecting their mental health and attrition in graduate programs. Yet we are only beginning to understand what contributes to graduate student anxiety. Biology Graduate Teaching Assistants (GTAs) have simultaneous roles as teachers, researchers, students, and employees, and factors associated with these tasks may contribute to anxieties in graduate school, particularly in relation to teaching and research responsibilities. To explore factors related to GTA teaching and research anxieties, and guided by social cognitive career theory, we interviewed 23 Biology GTAs at a research-intensive southeastern university. Thematic analysis of interview transcripts revealed five major factors related to GTA anxieties: **negative impact on self**, **negative impact on others**, **lack of self-efficacy**, **role tension, and personal anxieties.** Lack of self-efficacy was most prevalent for research anxieties, compared to teaching anxieties, where the impact on others (e.g. students) was most prevalent. In research contexts, GTAs with academic career aspirations expressed less anxiety about role tensions compared to GTAs with non-academic career goals. By investigating GTA anxieties, this work can inform professional development or mental health interventions for GTAs and encourage greater awareness and dialogue about mental health issues in academia.

## INTRODUCTION

Graduate students in the United States are experiencing increased levels of anxiety, affecting their overall mental health and attrition in graduate programs (Evans et al. 2018; Hish et al. 2019; Levecque et al. 2017; Nagy et al. 2019). A recent study found that one in three graduate students was depressed, a rate six times higher than the general public (Evans et al. 2018). Nagy et al. (2019) found high rates of mental health problems in biomedical doctoral students, with participants experiencing burnout, depression, and anxiety. Despite these reports of mental health issues in graduate students, we are only beginning to understand the contributing factors to graduate student mental health.

This study specifically focuses on anxiety (versus stress), which is defined as the state of anticipatory apprehension over possible deleterious happenings (Bandura 1988). Anxiety’s physiological responses are similar to those of stress: increased levels of cortisol, faster heartrate, dilated pupils, etc. However, with anxiety, these physical changes accompany general feelings of concern, tension, or worry about an anticipated event or outcome that may or may not actualize in the future; whereas stress may refer to the cognitive, emotional and biological reactions to specific and current life events (American Psychological Association 2020; Pekrun et al. 2007). Despite its anticipatory nature, intense anxiety can be debilitating to an individual as well as those around them. Though we recognize that some anxiety can sometimes be beneficial to an individual’s motivation or productivity (Cohen 2011; Pelton 2014), this study focuses on anxieties which are debilitating and negative for an individual’s well-being. By examining one aspect of mental health (anxiety) in our graduate students at great depth, this study aims to inform the factors that can support graduate student well-being.

### Identifying sources of anxiety of graduate students is important for improving their mental health

Our current understanding of the causes of general graduate student mental health issues, such as anxiety, are wide ranging—lack of advisor support, lack of social support, poor perception of employment prospects, or family/monetary concerns, to name a few (Devos et al. 2017; El-Ghoroury et al. 2012; Golde 2005; Hish et al. 2019; Levecque et al. 2017; Mousavi et al. 2018). For example, when Leveque et al. (2017) surveyed over 3,000 doctoral students in Belgium, they found that work-family conflict can exacerbate a graduate student’s risk of having or developing a common psychiatric disorder (Levecque et al. 2017). If these conditions persist for a graduate student, they can lead to attrition out of the program (Devos et al. 2017; Golde 2005).

Despite the growing number of studies that attempt to pinpoint the causes of graduate student mental health issues overall, we often fail to disambiguate specific parts of the graduate student experience that may have different impacts on anxiety, for example, graduate student roles related to teaching and research. Continuing to focus on the sources of anxiety for graduate student life overall, without considering the differential impacts of specific aspects of graduate student life, will make it harder to understand how to address graduate student anxiety. Therefore, this study takes a first step in comparing and contrasting the sources of anxiety related to teaching and research roles of graduate students.

### Anxiety related to teaching and research

Graduate students are important to universities for their work in teaching. Specifically, research universities rely on graduate students for instruction, especially for large enrollment classes, making the quality of their instruction important to undergraduate student success and retention. Biology Graduate Teaching Assistants (GTAs), for example, teach over 91% of freshman Biology labs and discussions nationally (Sundberg, Armstrong, and Wischusen 2005). The Longitudinal Study of Future STEM Scholars (LSFSS) found that nearly all (94.9%) of a 3,000 STEM PhD sample had taught undergraduates during their doctoral programs (Connolly et al. 2016). Within 5 years after graduation, almost half of those STEM doctoral graduates went on to teach in postsecondary institutions (Connolly et al. 2016), meaning they also continued to impact undergraduate instruction.

Graduate students, however, often find themselves teaching with little to no pedagogical professional development (Gardner and Jones 2011; Prieto and Scheel 2008), all while establishing research projects and integrating into departmental cultures. As a result, GTAs may experience a lack of confidence about their teaching (Pelton 2014; Prieto and Altmaier 1994; Reeves et al. 2018), resulting in teaching anxiety. Cho et al. (2011) identified a variety of GTA concerns that might produce anxiety, including class control, external evaluation, teaching tasks, student impact, holding dual roles, and time management. In a literature review exploring K-12 teacher anxiety, Coates & Thoresen (1976) identified sources of anxiety for beginning and experienced teachers. They found that beginning teachers were concerned with self, and experienced teachers were concerned with students’ educational growth and with personal teaching performance. Research in both K-12 and university contexts have found that teaching anxiety negatively impacts teaching behavior and performance (Cho et al. 2011; Coates and Thoresen 1976; Parsons 1973; Pelton 2014). Given university reliance on GTAs for teaching, factors that decrease instructional quality (such as teaching anxiety) may greatly influence the quality of undergraduate education at the institution.

Institutions not only rely on graduate student’s teaching for large enrollment courses, but also rely on their successful research output. With mounting pressure to “publish or perish,” graduate student productivity is critical to the successful functioning of large, research-driven universities (National Academies of Sciences Engineering and Medicine 2018). Despite this intense focus on research productivity, little research has been done on the sources of graduate student research anxiety and how it may impact their work. Though teaching anxiety and factors related to it have been measured and tested (Parsons 1973; Pelton 2014; Roach 2003), there are few studies which explicitly investigate graduate student research anxiety. For example, Brinkman & Hartsell-Gundy (2012) identified research anxiety related to a lack of information literacy skills, stating that “*while most graduate students were successful as undergraduates, many have never conducted a large-scale independent investigation, often resulting in heightened anxiety about their research projects*.” Therefore, to inform our work we had to draw from the closest relevant literature to GTA research anxiety which explores graduate student stressors, challenges, or concerns in their program broadly or studies focusing on specific anxieties related to subject matter (e.g. statistics, math), or tasks (e.g. the dissertation writing process) (Barry et al. 2018; Devos et al. 2017; El-Ghoroury et al. 2012; Golde 2005; Hish et al. 2019; Levecque et al. 2017; Mousavi et al. 2018).

Besides anxieties arising from teaching or research alone, another source of anxiety for GTAs may be related to tensions between teaching and research roles (Reid and Gardner 2020). Graduate students must learn how to balance multiple roles over their degree program (Kajfez and McNair 2014; Nicklin, Meachon, and McNall 2019; Winstone and Moore 2017). GTAs often teach undergraduate labs or discussions along with taking their own classes and conducting research (Gardner and Jones 2011; Prieto and Scheel 2008; Sundberg, Armstrong, and Wischusen 2005). Even when not pursuing a career in academia, many graduate students are required to teach as part of their assistantship, obliging them to balance roles they may or may not even want. Failure to successfully balance such tasks may exacerbate anxieties (Lane, Hardison, et al. 2019). Some graduate students are told directly not to spend time on teaching to reserve more time for research, creating external tension between roles (Lane, Skvoretz, et al. 2019; Reid and Gardner 2020).

Thus, in academia, anxiety related to a GTA’s teaching, research, or tension in balancing both may negatively impact graduate student well-being. By honing in on these specific aspects of the graduate student experience in great depth, this study can help to identify concrete targets to reduce teaching and research anxiety.

### Anxiety changes over time

Graduate school is a critical time for the socialization of graduate students into the professoriate (Austin, 2002), specifically into the academic norms, values, and beliefs that underlie teaching and research practice. Austin (2002) described socialization as “*an ongoing process, not the result of occasional events*” and as “*a two-way process where individuals both influence the organization and are influenced by it. It is a dynamic process in which the individual newcomer brings experiences, values, and ideas into the organization*.” Alignment of these values and beliefs takes time, suggesting that factors underlying graduate student anxiety may change over time as well.

Graduate programs represent distinct communities of practice into which GTAs enter and must integrate (Wenger 1998). The community has certain norms, values, and beliefs that graduate students need to recognize and adhere to over time. This means that their own norms, values, and beliefs will be under examination and perhaps need to be altered. Austin (2002) indicates factors such as a student’s locus of control and self-efficacy can impact this socialization process. In some cases, students do not integrate well into the community because they feel conflict between their own beliefs and that of the community. For example, they may want to spend time teaching, but are told to focus on research. They can explicitly fight these norms, or silently reject them. In all cases, this integration is an upheaval for the student and may cause anxiety as students seek to adjust to the new community. However, with integration may come a reduction of these anxieties. With this in mind, capturing the sources of teaching and research anxiety of Biology GTAs over time may provide insight into the extent to which anxiety changes and in what ways it might change.

Understanding how sources of GTA anxiety in teaching and research change over time can also help understand reasons for graduate student attrition and effective professional development (PD) for teaching and research at different stages of the degree program. For example, when examining doctoral attrition, Lovitts (2001) remarked that the decision by graduate students to leave a program is often not a one moment decision, but a collection of moments. Thus, there may be accumulating anxieties toward research or teaching over time that could be a factor in attrition that are currently unknown. This research would also help PD developers, coordinators, and instructors provide effective PD opportunities to graduate students at different points in their program. For example, if we found that experienced and novice GTAs have different sources of teaching anxieties, PD could cater differently to the needs of each to support wellness in each year.

### Theoretical framework: The Social Cognitive Career Theory (SCCT)

A graduate student’s career prospects may also inform their sources of teaching and research anxiety. Graduate student research productivity determines their career prospects, particularly if graduate students aim to pursue increasingly competitive academic appointments (Larson, Ghaffarzadegan, and Xue 2014). A study on biomedical doctoral students found that less than 20% of Ph.D.’s in the biological sciences moved into tenure-track academic positions within 5-6 years after receiving their degree (Fuhrmann et al. 2011), leading many graduate students to consider “non-traditional” career paths (Clair et al. 2017; Lindholm 2004), or solely teaching positions. One might infer that research anxiety in these aspiring future faculty would be relatively high. However, not all GTAs aspire to remain in academia as part of their future careers (Connolly, Lee, and Savoy 2018; Fuhrmann et al. 2011). The research anxiety of GTAs pursuing non-academic career aspirations may be relatively different from those pursuing academic positions. Using the Social Cognitive Career Theory (SCCT), we explore how a GTA’s career aspirations and the anxieties they hold towards research and teaching responsibilities may potentially influence each other.

The SCCT was developed to identify the cognitive and contextual variables which influence a person’s career interests and trajectory (Bandura 1993; Lent, Brown, and Hackett 1994) (**Figure 1**). For our study, we propose that these cognitive and contextual variables and the anxieties GTAs experience would influence each other, particularly anxieties related to their main roles in graduate school of teaching and research. The SCCT variables can help to identify where potential sources of anxieties may be rooted; for example, anxieties may be derived from an individual’s *learning experiences* (e.g. teaching in a classroom, conducting research in a lab), or from the cognitive variables of *self-efficacy* (e.g. “Can I teach / do research well?”) and *outcome expectations* (e.g. “What will happen if I teach / conduct research poorly?”), thereby impacting career-related development, choices, and performance in the future.

**Figure 1:**
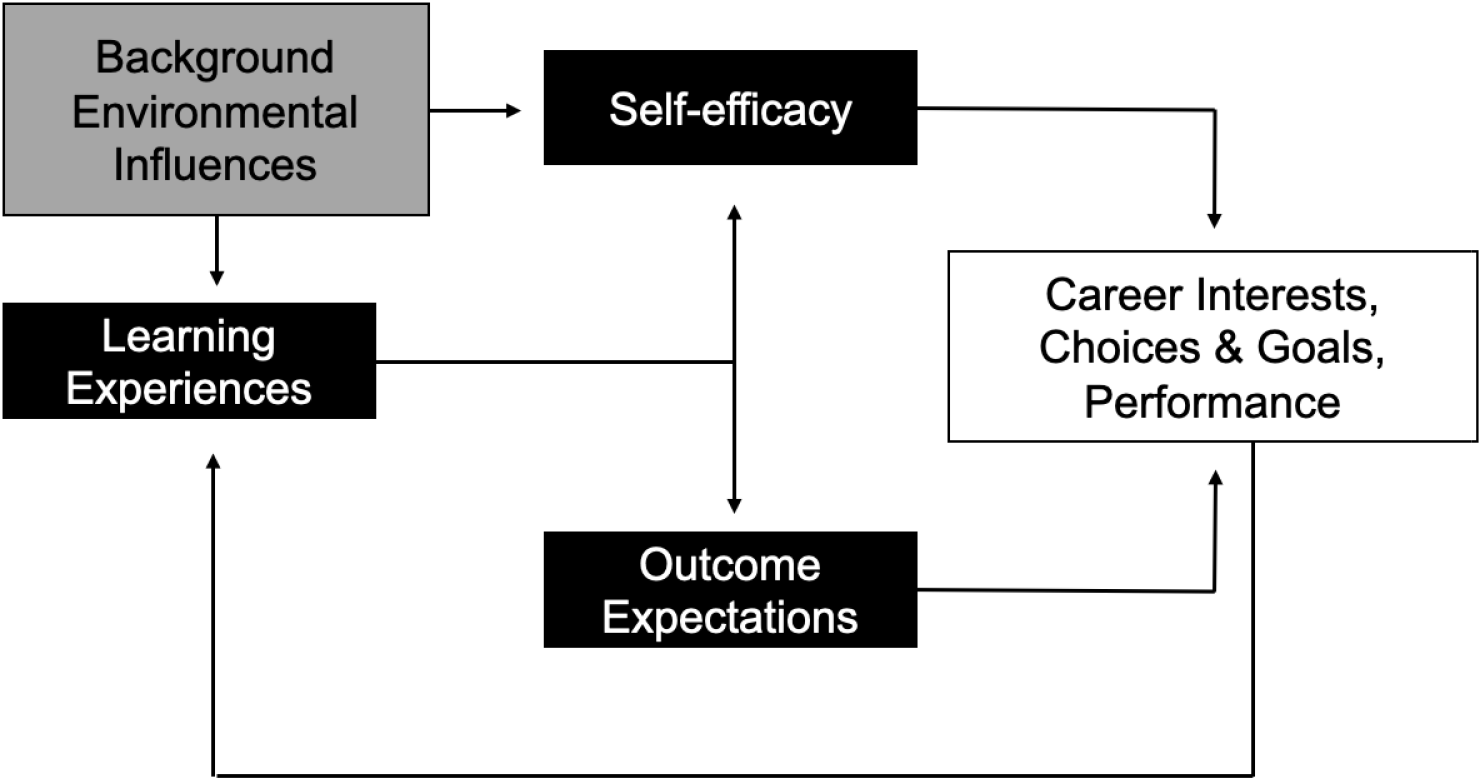
Theoretical model from the Social Cognitive Career Theory (SCCT), depicting the cognitive and contextual factors which influence career interest development. The SCCT aims to explain three interrelated aspects of career development: (1) how basic academic and career interests develop, (2) how educational and career choices are made, and (3) how academic and career success through performance is obtained. These three outcomes are displayed in the white box. The grey box indicates background or contextual components (e.g. demographics, teaching and research culture) and the black boxes are the main cognitive drivers that contribute to career development. We posit that the anxieties GTAs experience in their teaching and research roles may be related to these factors.

Guided by the literature, one of the main potential sources of anxiety can be found in the relationship between anxiety and the SCCT variable of self-efficacy. Self-efficacy is a critical component mediating anxiety and a key cognitive variable in career development and interest (Lent, Brown, and Hackett 1994, 2000). Self-efficacy is the belief or confidence in one’s ability to successfully carry out a specific task or course of action (Bandura 1988; Lent, Brown, and Hackett 2000) and has been studied in the GTA population (Connolly et al. 2016; DeChenne et al. 2015; DeChenne, Enochs, and Needham 2012; Hish et al. 2019; Reeves et al. 2016). Anxiety and self-efficacy often vary together in a feedback loop, with greater self-efficacy being related to less anxiety and vice versa (Bandura 1988). These two variables have downstream effects that impact the career interests, choices, and career performance of graduate students.

Cognitive variables such as self-efficacy and anxiety are dynamic; they change depending on a GTA’s learning experience and background influences, meaning self-efficacy and anxiety would be expected to change over the course of a graduate degree program (e.g. the impact of professional development on GTAs, see Connolly et al., 2016). As proposed by Bandura (1988), mastery experiences are one of the dominant pathways to building self-efficacy for a task, and thus reducing anxiety towards said task (Bandura 1988). For example, in a study modelling the relationship between stress, burnout, and depression in biomedical doctoral students, Hish et al. (2019) found that mastery of a skill mediated the relationship between stress and burnout, and between stress and depression (Hish et al. 2019). As graduate students build their skills in teaching and research, there should be a theoretical increase in self-efficacy and related decline in anxiety toward those tasks. Concomitantly, graduate students may also master balancing teaching and research roles (Austin and McDaniels 2006; Bucher and Stelling 1977), leading to reduced role tension and anxiety between teaching and research roles.

Using the SCCT model, these perceptions of self-efficacy and anxiety would further inform a GTA’s career aspirations, either by achieving or failing to achieve goals or tasks aligned with such aspirations. For example, evidence of task competence or enjoyment in teaching (e.g. positive classroom student evaluations) or research (e.g. successful, well-cited publication) would decrease anxiety related to those roles through validation of competency, high outcome expectations, and increased self-efficacy. The bolstering of self-efficacy and outcome expectations would positively encourage career pursuits where teaching and research are important roles (e.g. a research faculty position). While SCCT does not explicitly include the element of time in the framework, SCCT represents the *process* of career interest formation and the feedback loops in the model suggest the passage of time. As an example, a GTA may begin graduate school with no specific career interests, high self-efficacy for research and low self-efficacy for teaching, but find that over time they enjoy teaching, gain self-efficacy, and start to consider a future career in teaching. Conversely, a GTA who begins with a high self-efficacy for research and then has poor experiences with research tasks (e.g. too many rejected manuscripts), may then experience high anxiety with regard to research, declines in self-efficacy, and develop negative outcome expectations over time, leading to a disinterest in careers with a research component.

For the purposes of this study, we will be using the SCCT framework to explore the sources of teaching and research anxieties in Biology graduate teaching assistants (GTAs), including ideas about how or why these sources may change over time, and whether they differ with academic or non-academic career intentions of the GTA participants. Academic career aspirations were defined as careers as faculty or instructors at R1 and R2 institutions, liberal arts colleges, community colleges, and PUIs. These positions included having a research and teaching component, or simply with teaching responsibilities at the college-level. Non-academic career aspirations included seeking employment with government agencies, industry, non-profits, and elementary or high school teaching. Teaching-focused positions could fall into both categories (academic vs. non-academic careers), depending on the specific career aspiration. For example, an academic career goal to be an instructor at a liberal arts institution with some research would fall under the academic category. A career aspiration to be a K-12 teacher would be considered a non-academic career.

### Rationale and research questions

This study characterized the sources of anxieties related to teaching and research roles of graduate students at two points in time, and whether these sources of anxiety differ based on a GTA’s career aspiration for a group of GTAs in Biology at one institution. We explored the factor of time because of the potential for changes in sources of anxiety as graduate students adapted to the norms, values, and beliefs for their teaching and research roles in their academic learning community (Department). As the first study to investigate specific sources of teaching and research anxieties, we asked three main research questions:

1. What are the sources of anxiety for teaching and research roles in Biology GTAs?
2. Do these sources of teaching and research anxieties change over one year?
3. Do the sources of teaching and research anxieties differ based on GTA career aspirations?

Answering these questions will provide a more nuanced understanding of the sources of anxiety for graduate students, specifically for two common roles within a degree program. These findings can be then used to support graduate students’ well-being more globally.

## METHODS

### Study population

This study was approved by the University’s Institutional Review Board (IRB-16-03235-XP). Biology GTAs at this large research-intensive southeastern university were our study population. The GTAs were recruited from across the University’s Division of Biology through a listserv of graduate students from three departments and one program. As of Fall 2016, 211 graduate students in the Division of Biology were enrolled in a Masters or Ph.D. program, with 94% of graduate students seeking a Ph.D., and 55% identifying as female. We did not have access to institutional data in regard to other demographic characteristics of these students, so cannot provide information on ethnicity or other characteristics of the potential pool.

### Data collection

In October 2016, an online survey was deployed to Biology graduate students via the Qualtrics survey software. The e-mail targeted individuals who were either currently teaching or who had been a GTA previously. This survey collected quantitative data for another study, but was also used to recruit participants for this study. Specifically, at the end of the survey, participants were asked if they were “*interested in volunteering to participate in a brief follow-up interview?*” If they responded yes, they were prompted to enter their name and email so the researcher could contact them. Of the 89 Biology GTAs who completed the survey, 26% (n = 23) indicated that they would be interested in participating in follow-up interviews.

Interviews of graduate students were conducted twice over one year to collect in-depth perceptions of sources of teaching and research anxieties. All GTA demographics were collected using the survey disseminated in Fall 2016. We collected participants’ gender, ethnicity, citizenship status (international vs. domestic), teaching experience (number of semesters as a GTA), age, degree program, year in the graduate program, and department affiliation. We considered “experienced GTAs” as those having greater than or equal to 1 year of GTA experience, and “novice GTAs” as having less than 1 year of GTA experience as of the Fall 2016 interviews. GTAs in the interview pool were 70% female, and 74% white (**Table 1**). All interviews were conducted by the first author (M.C.M.). Each interview was approximately 60-90 minutes long. A small monetary compensation of $10 was offered to each graduate student for each interview, which was disseminated after the interview was complete.

**Table 1:**
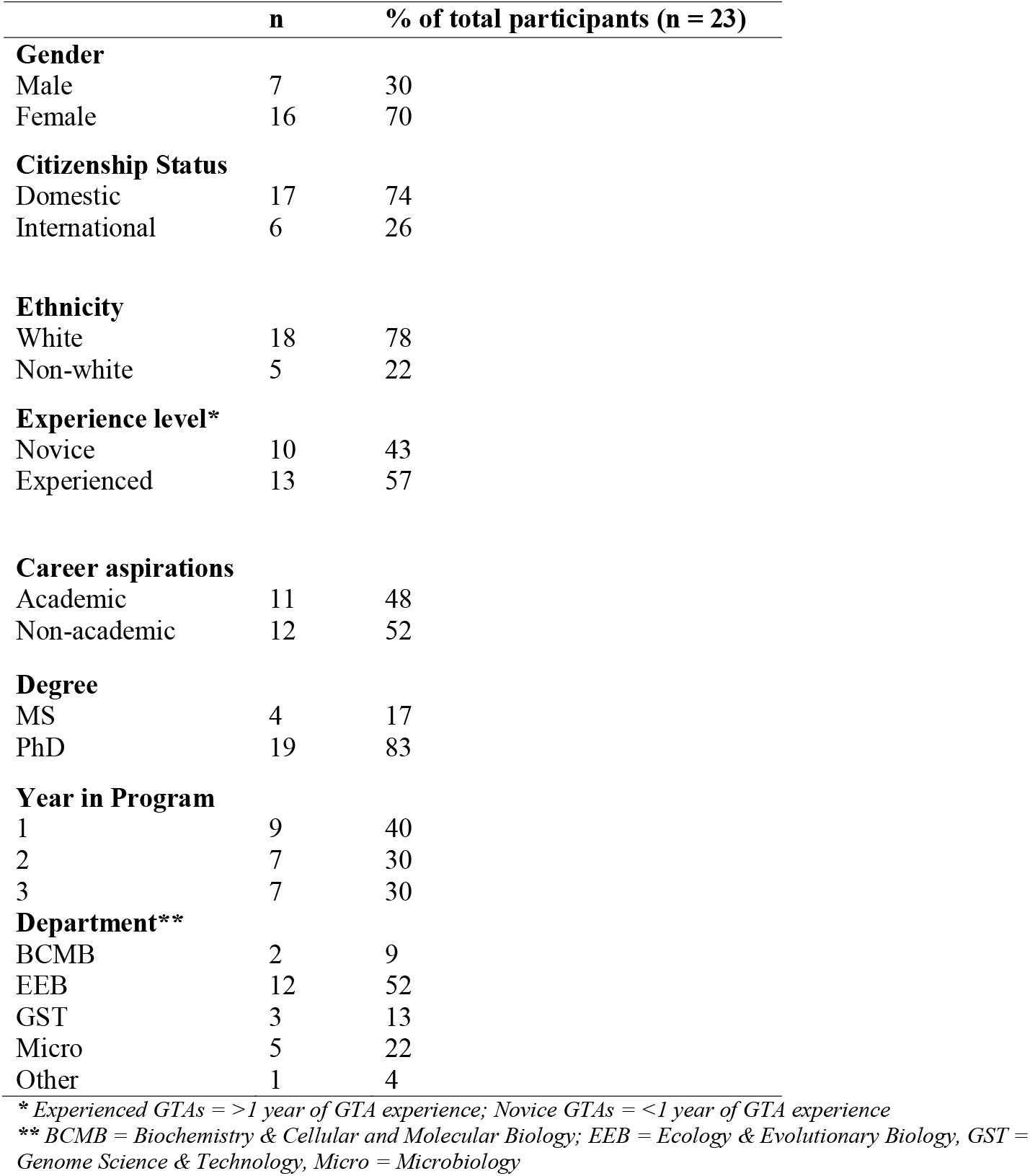
Demographics of the 23 Biology GTAs interviewed over 2016-2017.

|  | **n** | **% of total participants (n = 23)** |
| --- | --- | --- |
| **Gender** |  |  |
| Male | 7 | 30 |
| Female | 16 | 70 |
| **Citizenship Status** |  |  |
| Domestic | 17 | 74 |
| International | 6 | 26 |
| **Ethnicity** |  |  |
| White | 18 | 78 |
| Non-white | 5 | 22 |
| **Experience level\*** |  |  |
| Novice | 10 | 43 |
| Experienced | 13 | 57 |
| **Career aspirations** |  |  |
| Academic | 11 | 48 |
| Non-academic | 12 | 52 |
| **Degree** |  |  |
| MS | 4 | 17 |
| PhD | 19 | 83 |
| **Year in Program** |  |  |
| 1 | 9 | 40 |
| 2 | 7 | 30 |
| 3 | 7 | 30 |
| **Department\*\*** |  |  |
| BCMB | 2 | 9 |
| EEB | 12 | 52 |
| GST | 3 | 13 |
| Micro | 5 | 22 |
| Other | 1 | 4 |

\* Experienced GTAs = >1 year of GTA experience; Novice GTAs = <1 year of GTA experience
\*\* BCMB = Biochemistry & Cellular and Molecular Biology; EEB = Ecology & Evolutionary Biology, GST = Genome Science & Technology, Micro = Microbiology

### Biology GTA Interviews

Interviews were conducted with 23 Biology GTAs in Fall 2016, and again with the same sample of participants in Fall 2017 using the same interview protocol with the addition of a few retrospective questions. Interview questions probed four main topics: (1) a GTA’s perception of their experience level, knowledge of teaching, and effectiveness in teaching; (2) their teaching and research anxieties; (3) coping strategies enacted; and (4) their professional identity/career aspirations as teachers and researchers. For this study, qualitative analysis was conducted on participant responses to the second and fourth topics only in order to answer our research questions related to sources of teaching and research anxiety each year and how it may relate to career aspirations.

To probe the second topic, card sorts were used as a tool to guide conversations about factors that may be related to teaching and research anxieties. Card sort activities make abstract concepts more tangible for participants, especially as interactive, object-based aids to qualitative interviews (Conrad and Tucker 2019). It is a method employed to examine cognitive structures and processes (Spradley 1979; Weller and Romney 1988), and encourage the individual to express internal cognitions and schemas (Blake et al. 2007). For these interviews, one set of cards contained hypothesized factors or topics that may be related to teaching anxieties (e.g. student behavior, grading, etc.) and the other set contained hypothesized factors that may be related to research anxieties (e.g. writing grants, data analysis, etc.). Blank cards were also available for participants to write their own factors not captured by the existing cards.

The topics listed on the cards emerged from the collective experience of 2 faculty members (both of whom supervise graduate students in research and teaching roles), one Biology education postdoctoral research associate, and 2 Biology graduate students, all trained in biology and biology education research (including authors M.C.M. and E.S.). Together, we reflected on our own graduate school experiences and listed the things that caused us anxiety and then further thought about the factors causing anxieties in the graduate students in our department and the Division, specifically in teaching and research roles. Some of these ideas were related to general topics found in the literature pertaining to stress in graduate school (e.g. advisor relationship) and teaching (e.g. not knowing the content). However, our efforts went beyond what was in the literature, given the lack of empirical work on specific factors related to research anxieties in graduate students.

To prevent these pre-determined card topics from constraining participant responses during the interview, before reading any of the cards, we first asked participants to list any teaching and research anxieties on a sheet of paper (Supplemental Material). If the card sort topics failed to capture a source of anxiety for a participant, it would be captured in this initial anxiety listing. None of the participants indicated a total absence of anxiety for either teaching or research. Participants were then asked to read the cards from one set (teaching or research), and pick the topics that caused them anxiety for teaching and research, respectively. They then explained what about the topic on the card made them anxious, and the perceived impact of this anxiety on their teaching or research. At this point, participants would sometimes discuss the anxiety factors they wrote on the blank sheet of paper, which were not captured in the existing set of cards. Among the 23 participants, 9 Biology GTAs added new topics (3 topics for teaching, 7 topics for research). These cards were not added to subsequent interviews because we wanted to maintain consistency in the card sets and there was not enough time between interviews to amend the IRB interview protocol. Interviews were audio-recorded and transcribed in order to conduct thematic analysis (Boyatzis 1998; Braun and Clarke 2006).

### Data analysis

To identify themes and categories regarding sources of teaching and research anxiety of the participants, thematic analysis of interview transcripts was conducted first using the Fall 2016 interview data. This process sought to capture the experience and perceptions about factors related to teaching and research anxiety of GTAs (Charmaz 2006; Saldaña 2012; Strauss and Corbin 2008). Braun and Clark (2006) define *thematic analysis* as “*a method for identifying, analyzing, and reporting patterns (themes) within the data. It minimally organizes and describes your data set in detail*,” where the research topic informs the interpretation of the text (p. 79). The process was inductive, wherein the researchers identified major codes, categories, and themes without any predetermined codebook (Saldaña 2012). The researchers were aware of the theoretical framework of SCCT throughout the work, which likely guided some of the final themes that were created; however, the underlying codes and ideas were guided by the data and not explicitly by aspects of SCCT.

It is important to note that the role of the card topic was to generate a participant *explanation* for a source of their anxiety and not to necessarily record the topic as the source of anxiety. In other words, the card was a prompt for a discussion about anxiety with participants, with the coding focusing on the explanation for why they chose that card. For example, two GTAs may have picked the card “grading” as something which prompted anxiety for them. However, the first GTA articulated that being able to grade fairly caused them anxiety, while the second GTA explained that their anxiety was based on how much time grading takes away from research responsibilities (**Figure 2**). In this instance both GTAs chose the same card, however, the coding was solely based on the elaboration of the cause for anxiety. Our full interview protocol and cards are available in the **Supplemental Material**.

**Figure 2:**
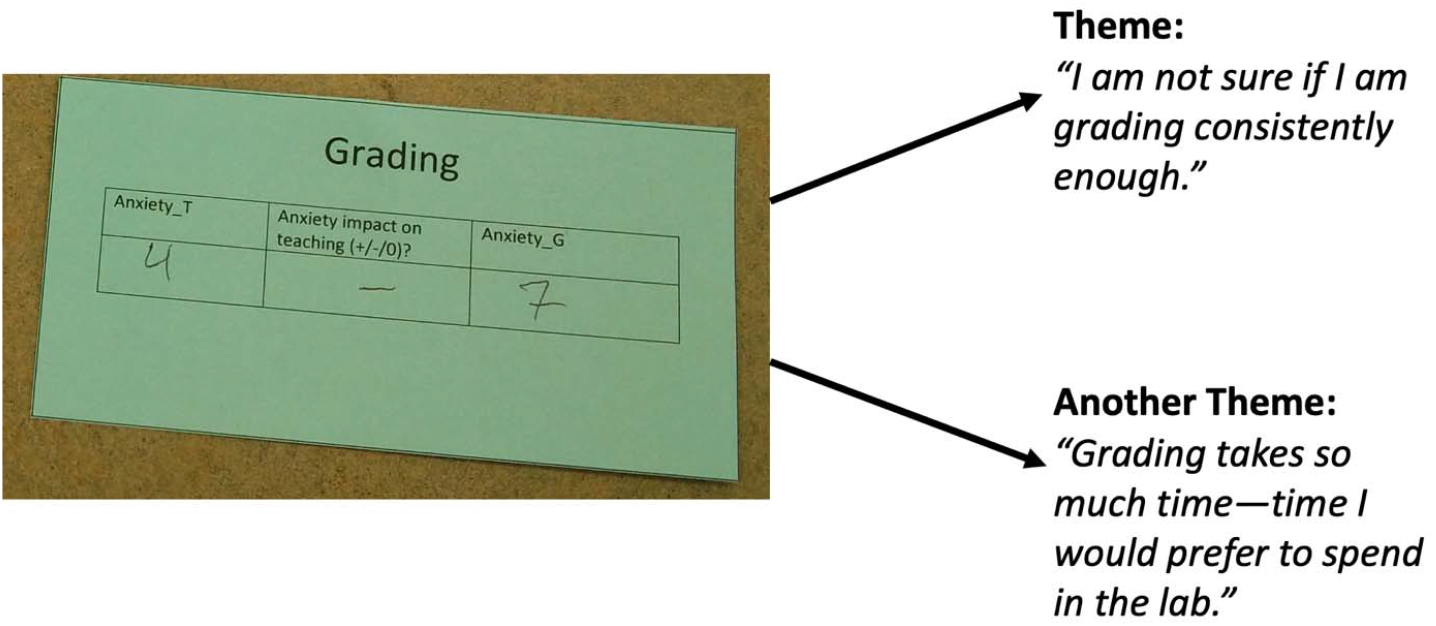
Example of a card provided to Biology GTA participants to probe teaching anxiety. Qualitative coding examined the explanation of why participants chose the card, which could sometimes differ among participants. In this example, two teaching anxiety codes emerged based on what participants said caused them anxiety related to grading.

The initial codebook was created by the first author after reading the Fall 2016 interviews. A method of constant comparison was then followed by the first author (primary coder) and two undergraduate research assistants (RAs, including co-author K.P.) to analyze interview transcripts and refine the codebook. All three coders began by independently coding the same set of six randomly selected interview transcripts from 2016, then came together to determine the percentage of code agreement. Coding units were participant responses to each anxiety card chosen for teaching or research. If there was disagreement lower than 80%, coders would re-examine the codebook, refine definitions of codes and their broader themes, and re-code a new set of transcripts. As new codes emerged, or new understandings developed, the codebook was revised and previous interview transcripts were re-analyzed to look for the new codes (Glaser and Strauss 1967; Schwandt 2007). From these descriptions and codes, major themes were then identified in the codebook. A major theme provided explanation for distinct aspects of the overarching finding. The codebook themes and descriptions were reviewed by 2 faculty members (both trained biologists and biology education researchers), one biology education post-doctoral researcher, and 2 biology graduate students to ensure codebook clarity.

Three iterations of this method were conducted among coders to reach a minimum agreement of 80% consensus of themes (Landis and Koch, 1977). For this study, a coding unit was a participant’s explanation for each card they choose, and could encompass more than one code. Therefore, we chose to calculate a percentage of agreement as a measure of inter-rater reliability (IRR) since methods like Cohen’s Kappa assume one code per coding unit. IRR is a measure of agreement among raters; greater consensus among raters indicates higher reliability of the codes (LeBreton and Senter 2008; McHugh 2012). IRR was calculated by percent agreement among the first author and 2 other undergraduate research assistants after each coding iteration by:

1. summing how many times coder 1, 2, and 3 agreed per coding unit;
2. then determining the percentage of agreement between coders among each interview.

For example, if there were 4 coding units to be coded, and coder 1 and 3 agreed on all the codes, but coder 2 disagreed on only one of the units, then the IRR percent calculation would be as follows: 3/3 + 3/3 + 3/3 + 2/3 = 92% agreement, since 3 of 3 coders agreed on 3 interview segments, while only 2 coders agreed on the last interview segment. After the last coding iteration, among 3 coders, 98% agreement was calculated across 3 different interview transcripts from Fall 2016.

The codebook which inductively emerged (Merriam and Tisdell 2016) from the analysis of Fall 2016 interviews was used to code subsequent interviews conducted on the same group of GTAs in Fall 2017. The first author finished coding the remaining Fall 2016 interviews, and the two RAs coded interviews from Fall 2017. We verified that data saturation was met by the end of the 2016 interviews because we were hearing similar ideas from participants. We verified this via coding because no new themes emerged between the 2016 and 2017 interview analysis. All coding was recorded and conducted in Microsoft Excel, with each interviewee having one Excel Sheet and each coding unit occupying an independent row.

To answer the first research question, thematic analysis of interviews identified major themes that encapsulated the sources of teaching and research anxiety that the graduate students expressed. To summarize and compare the teaching and research themes, we tallied the presence and absence of each theme for each Biology GTA participant from the 2016 and 2017 interviews, respectively. If a transcript contained multiple instances of codes related to the same theme for the same participant, we only counted the theme once for that participant. For example, if a participant indicated they were anxious conducting a new statistical analysis for their research and also mentioned anxiety about properly collecting the data, we would only designate that participant as having anxiety related to lack of self-efficacy once. After tallying the emergence of each theme among GTA participants, we calculated a total percentage of themes among the participants. For instance, if 19 participants indicated anxiety related to lack of self-efficacy, the percent emergence of that theme in the population would be (19/23) × 100 = 83%. The total percent emergence of each theme for research and teaching anxiety was then compared.

To answer the second research question, examining how the sources of teaching and research anxieties compared from 2016 to 2017, we examined the themes identified for each participant in 2016 and 2017 and then determined if those themes were kept (present each year), added (present only in 2017), resolved (present only in 2016), or were never present (not present in either year). Once we determined the changes over one year for each participant, we then tallied how many participants were in the four categories of change (or no change). Lastly, to answer the third research question, we compared percent emergence of themes between the academic vs. non-academic career choices of each participant. Pseudonyms, aligned with participant’s gender and ethnicity, are used throughout the results section.

### Methodological limitations

Our study is bounded by the 23 Biology GTAs we interviewed, and their experiences at a particular time (2016-2017). We cannot broadly generalize or claim that these themes would be found in other graduate student populations. This study was also bounded by the card topics given to the Biology GTA participants. Though participants were given the opportunity to articulate anxieties they experienced in teaching and research before any cards were presented to them, it is possible the card topics constrained some of their responses. However, as the first qualitative study to examine GTAs’ anxieties pertaining to research and teaching roles, we hope academic scholars and practitioners responsible for the training of GTAs consider what these GTAs have indicated in our Results as a starting point for their own contexts. As Merriam (1995) argues, the purpose of qualitative research is not for generalization but to capture the essence and phenomenon of what that bounded population is experiencing.

Secondly, since this is a sensitive, often stigmatized topic, we recognize that our sample represents individuals who were comfortable sharing their anxieties, and likely does not represent the whole GTA population. There are existing barriers that individuals face when talking about or reporting anxiety and mental health. Reluctance to seek help and divulge these mental health issues are often attributed to the fear of stigma or expected negative impacts on career aspirations, especially with men (Evans et al. 2018; OECD 2014).

## RESULTS

### Five themes characterized teaching and research anxieties among Biology GTAs, but proportions differed by teaching or research role

The interviews revealed five major themes related to sources of GTA teaching and research anxieties across 2016 and 2017 (**Table 2; Figure 3**). The first theme, **perception of others and its negative impact on self,** represent factors that cause anxiety related to how others perceive your work and judge you as an effective teacher or researcher (**Table 2a**). This theme is concerned with perceptions of the personal consequences (e.g. the GTA’s reputation) that may arise as a result of their actions or behaviors. In research, the perceptions of their main research advisor, other faculty, and peers were of most concern to participants. For example, Mark, a Master’s student pursuing a career in government, explained how he was anxious about being perceived poorly by his research advisor and removed from the lab:

> *“…Yeah, I think for the longest time I had been under this perception where if I mess up I’m gonna get kicked out. Which is something that I find to be irrational because I don’t think [my PI] would ever do that. But, I also generally take the assumption that I’m not doing a good job.” – Mark (Fall 2016)*

**Figure 3:**
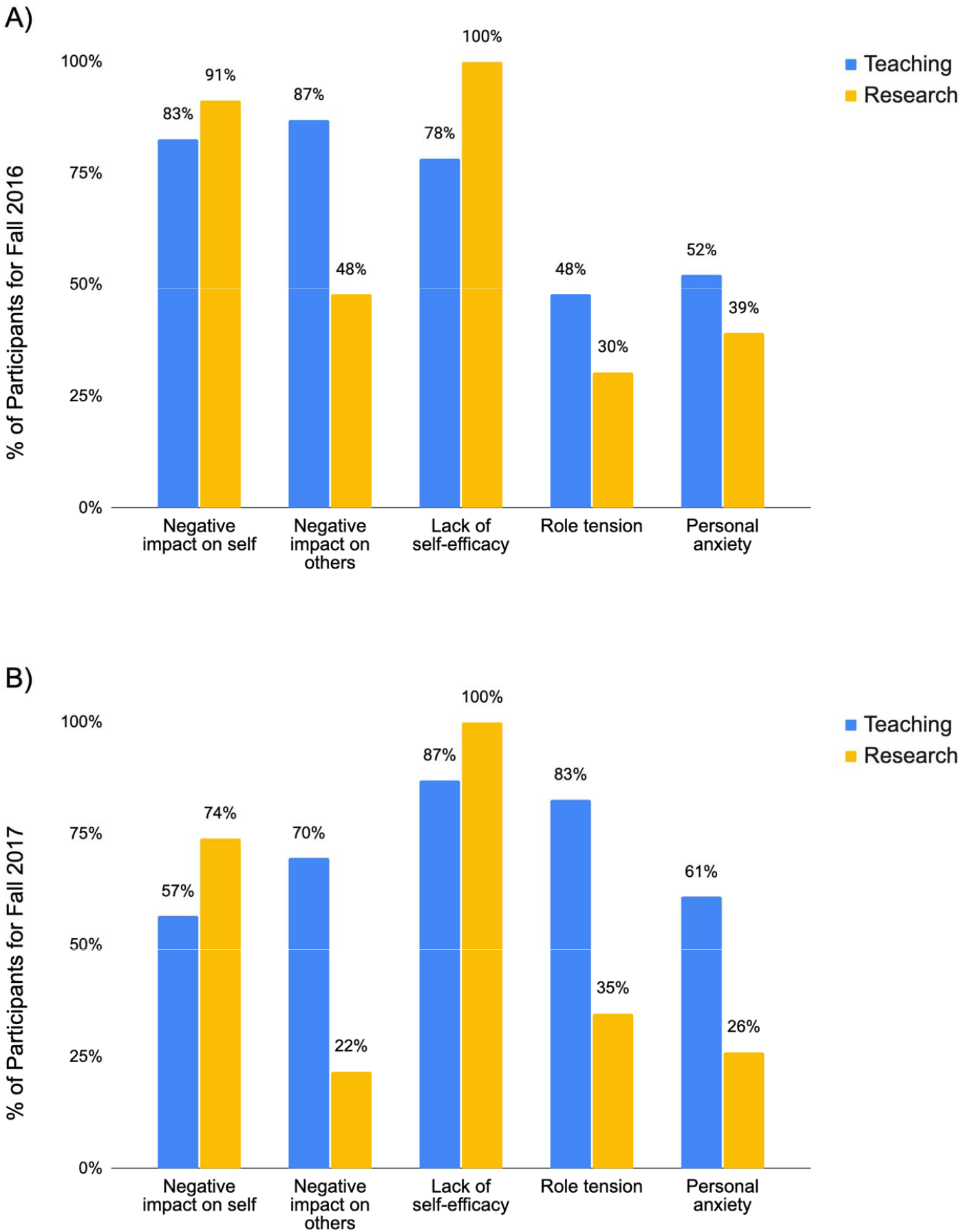
The percentage of participants (n=23) in which each anxiety theme emerged for teaching (blue) or research (yellow), in (**A**) Fall 2016 and (**B**) Fall 2017.

**Table 2:** Major themes (a through e) related to teaching and research anxieties which emerged from interview participants (n=23) from 2016 and 2017. Theme definition and an illustrative quote for teaching and research are included.

| Theme | Definition | Teaching Anxiety Quote | Research Anxiety Quote |
| --- | --- | --- | --- |
| **a) Negative impact on self**<br>(Theme 1) | Anxiety and fear related to how others perceive your work and judge you as an effective teacher/researcher. Anxiety related to the personal consequences of being unable to fulfill the expectations of oneself or others in teaching/research and its reflection on you. This theme pertains to how your behavior may change another person's thoughts of you and hurt you and your reputation. This theme is "you-oriented". | *"...even though I am more comfortable not knowing things, there is still the thought of 'Oh, well this is something you should know, and they're going to think that you don't know anything, and you can't teach them if you don't know anything.'"*<br>- Reagan | *"Normally, I don't care what people think of me, but **when it's with my peers, I'm more worried or concerned that they perceive me as someone who is here and serious about doing their research.** I'm always afraid of how I look like when I can't answer a question or mess up on a protocol. It could make me look like I'm not serious about this."*<br>- Emily |
| **b) Negative impact on others**<br>(Theme 2) | Anxiety of the consequences and impact on others if you are a poor or incompetent teacher/researcher. Participants expressing this anxiety are concerned about how one's actions (or lack thereof) may hurt another person (e.g. student, advisor). This anxiety is distinct from the previous category because it is rooted in how your behavior can hurt another. This theme is "others-oriented". | ***"I know that the decisions I make about grading, at least for most students, will affect them and might really make them sad or discourage them, and you try really hard not to do that, but you also can't just be a pushover about everything. Otherwise, you're not helping the students at all."***<br>-Julia | S: *"I think I just get worried. If I don't know [how to answer a question posed by my advisor], **I'll be a disappointment** kind of thing and that's one of my worst fears."*<br>I: *"That people would be generally disappointed or like?"*<br>S: *"Like **specifically my advisor and faculty committee people.**"*<br>-Samantha |
| **c) Lack of self-efficacy**<br>(Theme 3) | Anxiety related to the absence of a skill or control in situations of teaching/research. This anxiety is also related to inexperience, uncertainty, and lack of confidence in carrying out a task/handling a situation successfully. Also related to lack of | ***"Sometimes, I don't recognize right away what's the best way I should redirect, or what's the best [teaching] practice I should use. So, when the class redirects and I feel like I lose control a little bit,*** | *"I started coming up with my own experiment/ideas. So, it's **uncharted territory** for me and that gives me anxiety. I'm thinking about **'Has anyone done this before? Is there a better way*** |
|  | autonomy or knowledge in how to handle a situation or complete a task. | *that can stress me out because I feel like I'm trying to quickly redirect and the students pick up on it."*<br>-Lauren | *to do this? Is this the right way?"*<br>-Emily |
| **d) Role tension**<br>(Theme 4) | Anxiety related to lack of time to spend on other perceived priorities, particularly research, course work, and personal life. This theme is typically in the context of teaching taking away too much time from research. Tension in the roles and responsibilities a GTA has in their program. | *"I just feel in the course I'm currently [teaching], I spend way too much time grading every week, and it takes away from my research...and I'm just grading really crappy worksheets that aren't really helpful to the students in the first place, and their answers aren't very good because the questions aren't. It's a lot of time spent on something that's no good for me or for the students."*<br>-Cathy | *"...pretty much everything that I'm doing, I'm thinking I should be doing research all of the time. If I am even doing errands for the lab, I'm thinking: 'I should shove this on someone else, so I can get back to doing my research'"*<br>-Eric |
| **e) Personal anxiety**<br>(Theme 5) | Anxiety and stress related to personal worries or problems. This is often related to lack of finances, paying off loans, politics, religion, etc. This theme often comes up when the "Living off my stipend" card is chosen by participants. | S: *"I like the actual dissection in our [TA] meetings, but it sucks 'cause it's on Sundays. Six hours on Sundays...so it's hard to schedule around that. My girlfriend is 4 hours away, so I usually go on the weekend [to see her] and that takes the option out of the way. Just being on the weekend kind of sucks."*<br>-Jack | *"So, as a grad student, we all wear a lot of hats ..., but I still have a home life - I've got a husband at home that would like me to spend some time with him. I still do the thing where we're sitting on the couch watching TV, but I have my computer open doing work. So, work/life balance is always a thing I struggled with, and he realizes this. I'm driving down to a conference with my husband, but he sees this as a vacation - I see it as I've got work from 8am-6pm."* |
|  |  |  | -Laretta |

In teaching contexts, the perceptions of students, instructors, and teaching observers were those in which participants were often most concerned about. Perceived consequences of negative perceptions were a loss of student/classroom control or respect in the classroom or lack of respect or trust in teaching effectively from instructors. To illustrate this theme, Kaitlyn, a novice GTA also pursuing work as a government researcher, spoke about how she was anxious that her students would lose their respect for her if she was unable to answer a student’s question:

> *“…being unable to answer the question might affect the students’ perception of you…. If their opinion of you becomes lower, they‘re not going to respect you as much as a teacher.” – Kaitlyn (Fall 2017)*

The percentage of GTAs who articulated this theme was higher for anxiety related to research in both Fall of 2016 and 2017 (91% and 74%, respectively) than for teaching (83% and 57%, respectively) (**Figure 3**).

Similar to the first theme, the second theme which emerged among Biology GTAs was **the perception of others and its negative impact on others,** or anxiety in which the GTA is concerned with how their actions or behavior could harm their students, research advisors, or coworkers, rather than themselves (**Table 2b**). Compared to the first theme, which is more inward or self-facing (“how can this impact me”), this theme is focused more on consequences which are outward or others-facing (“how can this impact those around me”). In research, this would be related to anxieties about how GTAs perceived actions or behaviors would impact their main research advisor, their lab mates, colleagues, or even the wider community. For example, Sunny, an international doctoral student with industry and entrepreneurial career aspirations, indicated an anxiety in carrying out meaningful research to help people:

> *“[I want] to make [my research] useful. To prove this will work. This way can help people.” – Sunny (Fall, 2017)*

In teaching contexts, the major narrative for this theme involved a GTA’s fear of failing to teach a concept competently to their students, thus affecting their students’ course grade. For example, Jose, an international student and novice GTA pursuing academic career goals as a research professor, explained how he “feels bad” when he fails to properly teach a fundamental concept to his students:

> *“…not being as competent as I would like now, affecting them long-term in the course. Maybe I’m not teaching this part very well, where everything builds upon it. Like the first two-thirds of the course will be based on the first week and a half of material…. Then, you’re gonna have problems throughout…Maybe you lose ‘em, maybe you confuse ‘em…that’s even worse.” – Jose (Fall 2017)*

This theme occurred more frequently when describing teaching anxieties (87% Fall 2016; 70% Fall 2017) than research anxieties (48% Fall 2016; 22% Fall 2017).

The third theme – anxiety rooted in **lack of self-efficacy** – was one of the most predominant themes (**Table 2c**). This theme is characterized by feelings of uncertainty or inexperience towards a teaching or research task, and doubts of self-efficacy, autonomy, or control at a given teaching/research situation. Participants articulated how the lack of self-efficacy are related to feelings of “no control” in handling novel or unknown tasks in teaching or research. This theme commonly appeared when the GTA was required to do a task for the first time or perceived difficulty when encountering certain tasks or situations in the future. In research, some of these tasks or situations would be conducting data analysis or data collection, writing for publication, or graduating and finding a job afterwards. For example, Kayla, an international Master’s student aspiring to be a professor, described how she lacked guidance for her field work, therefore she questioned her ability to perform her research properly:

> *“When I was doing field work this season, I usually have to do it by myself. I didn’t have anyone helping me, and there are so many times where I was like ‘I don’t know what this is‘, but I had to think quickly there, and while I’m doing things, I’m like ‘I hope I’m doing this right.’” – Kayla (Fall 2017)*

For teaching, a few instances of the tasks or situations which gave rise to anxiety in Biology GTAs would be in teaching a new course or new material, grading fairly, assessing student learning accurately, or being able to “reach” and positively impact their students. For example, Eric, a doctoral student with many semesters of teaching experience and non-academic career goals in science policy, indicated anxiety related to grading:

> *“I have this anxiety that I’ve started to just drift and I’m grading more lenient or harsh—especially with really subjective assignments…It’s hard to [grade consistently] and also grading fairly in general sometimes.” – Eric (Fall 2017)*

Similarly, Julia, a novice GTA primarily pursuing teaching-focused career goals outside of academia, also expressed a lack of self-efficacy related to how to deal with a classroom emergency:

> *“I actually don’t feel very prepared for emergency like what happens if there’s a shooter or something. They don’t really teach us that so that would be kind of scary.” – Julia (Fall 2017)*

This theme was consistently present for research for 100% of the GTAs from Fall 2016 and 2017. While not as prevalent, a large portion of the GTAs mentioned this theme for teaching (78% Fall 2016; 87% Fall 2017).

The fourth theme was **role tension** (**Table 2d**). This anxiety is based on time management issues, particularly concerns where time spent teaching or on teaching tasks take away from other priorities. This theme arose within two different contexts 1) tension between teaching and research responsibilities; and 2) tension between graduate school responsibilities and personal life. This theme was most often articulated when participants perceived teaching responsibilities as a lesser priority, and teaching was taking too much time away from research responsibilities. GTA Rebecca, a doctoral student pursuing career goals in full-time research outside of academia, explained, “*I’m not in grad school to teach. I’m in grad school to complete a higher education degree*.” This tension emerged mostly when GTAs were describing anxieties related to teaching, but not as often within research contexts. Again, illustrating the typical context this theme appears in, Sunny, lamented spending too much time on teaching, perceiving research as the greater priority in graduate school:

> *“I think research should be always the priority, but actually, I spend a lot of time in teaching.” – Sunny (Fall 2017)*

The second context for this theme was when graduate school responsibilities interfered with life outside of being a student. In other words, how being a student took time away from being a “normal person,” or having a “normal personal life.” Anika, an international doctoral student aspiring towards a non-academic career in industry, explained her anxiety over “juggling” all of her priorities as a graduate student, while maintaining a personal life:

> *“…you have a life outside school, it’s not like you can dedicate twenty hours to grad school. So, you want to make sure that you meet your friends and have a social life, but also do research, also write, and…make your boss happy. So, it’s a lot of juggling.” – Anika (Fall 2017)*

Similarly, Hannah, another doctoral student pursuing work in the government or industry, succinctly said:

> *“I have to spend a lot of time preparing for teaching. That takes away from time doing other activities like research or life.” – Hannah (Fall 2017)*

For Fall 2016, this theme was relatively moderate for GTAs in teaching (48%) and research (30%); however, in Fall 2017, this theme increased dramatically for teaching (83%) and increased only slightly for research (35%).

The final theme which emerged related to **personal anxieties** (**Table 2e**). This theme was distinct from the previous theme in that it lacks the time element. These anxieties were personal issues that arose in conjunction with being a graduate student (e.g. finances, politics, religion, familial issues, etc.). Personal anxieties were categorized under teaching or research contexts depending on when the Biology GTA articulated such anxieties. For example, many Biology participants chose the “Living off your stipend” card as a point of personal anxiety for teaching, explaining that being a GTA did not pay enough money. Some participants chose the card “Living off your stipend” if they perceived it also contributed to their research anxiety. In this example from Laretta, an experienced GTA with career aspirations to be a professor in academia, she expressed anxiety about the future changes to her family and life that will come post-graduation, including issues of money, geography, and family:

> *“I think the future is always a scary place…not only with the current funding climate, but trying to figure out where I should transplant my family to-I just got married, and we have so many pets at the house. We just bought a house last year, so what am I supposed to do? Am I supposed to leave my husband and pets at home-do post doc for 2 years and come back? Or bring everyone? Should I rent or sell my house? I don’t know. I don’t think that it would be difficult for me to find a post doc position, it would be difficult for me to figure out where and how to deal with it.” - Laretta (Fall 2017)*

When listing teaching anxieties in Fall 2017, Lauren, a novice GTA with career aspirations to be a researcher outside of academia, articulated anxiety related to recent federal policies that would potentially impact her teaching stipend:

> *“Well I think the whole tax cuts and job act has got me stressed recently and it’s just something that I’m thinking a lot about. It’s been on the forefront of my mind unlike in previous semesters. I’m just very conscious about it now on where I want to be financially when I finish and trying to think about what my future is going to look like…Just having money saved away, being able to support myself in an emergency and spending my money wisely is something that I’m a bit stressed about.” – Lauren (Fall 2017)*

For Fall 2016, 52% of participants expressed this theme in teaching and 39% for research. In Fall 2017, 61% of participants indicated it for teaching and 25% for research.

### Over time, there were changes in sources of anxieties for teaching and research for GTAs

One year after the initial interviews, changes in participants’ sources of anxieties differed between teaching and research contexts (**Figure 4**). In teaching, anxiety related to negative impact on self, negative impact on others, lack of self-efficacy, and role tension were most often kept over time versus resolved, added, or never had. Negative impact on self, however, was kept as an anxiety in the participants almost as often as it was resolved in 2017 (**Figure 4a**). The personal anxiety theme had relatively even numbers of participants keeping, resolving, adding, or never having it. In research contexts, anxiety related to negative impact on self and lack of self-efficacy were kept by most participants (**Figure 4b)**. Anxiety related to negative impact on others, role tension, and personal anxieties were most often resolved or never had. Comparing anxieties in teaching and research over time, negative impact on others, role tension, and personal anxiety, remained present in teaching contexts, either being kept or added, more prominently than in research. The details of how each Biology GTA participant changed in their anxiety in teaching and research from 2016 to 2017, as well as individual demographic information, can be found in the Supplemental Materials (**Supplemental Tables 1 and 2**).

**Figure 4:**
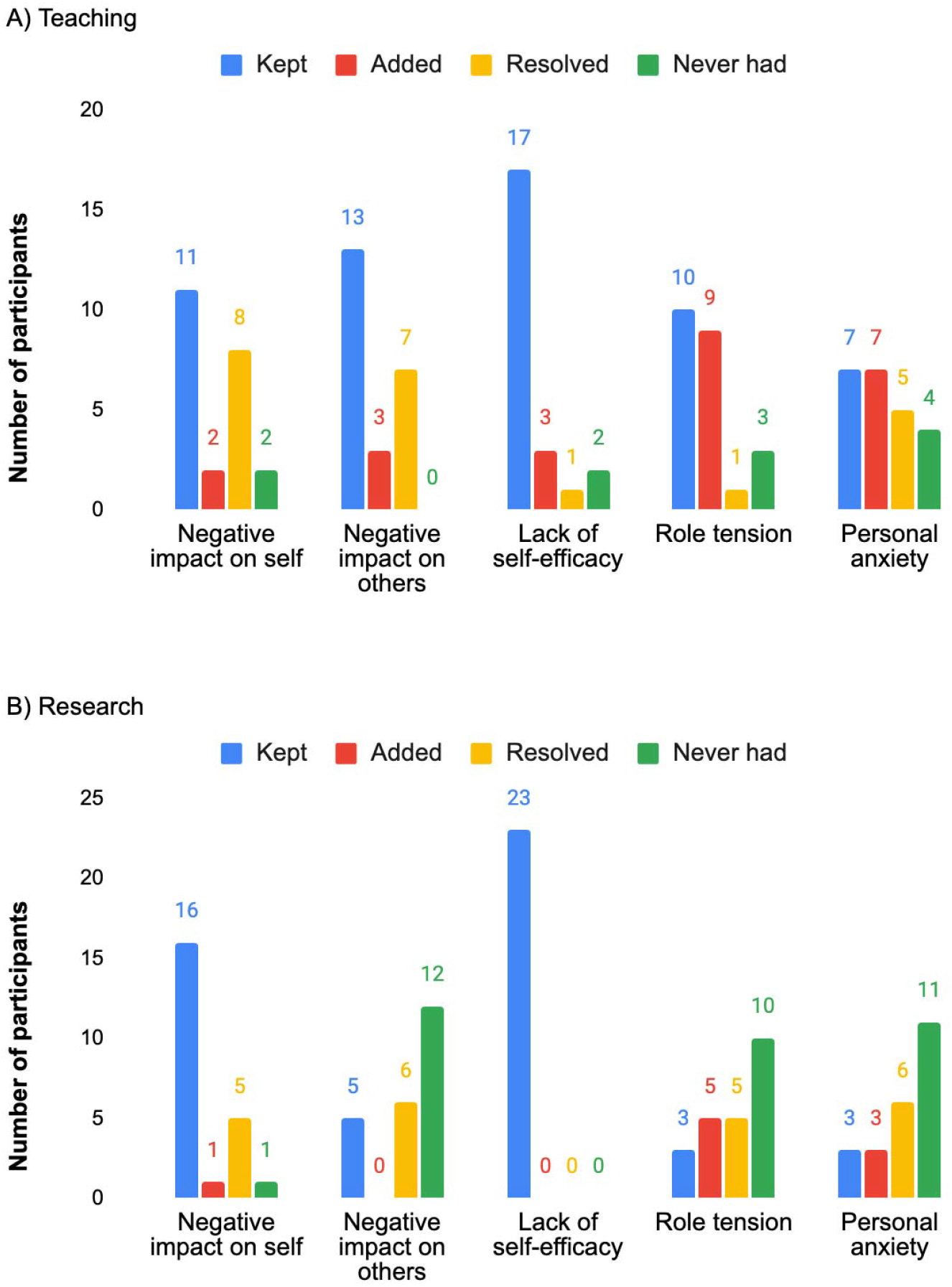
The number of Biology GTA participants which either kept, resolved, added, or never had that particular cause of anxiety from 2016 to 2017 in **A)** teaching and **B)** research contexts (n=23). The most common cause of anxiety which was maintained in both teaching and research was lack of self-efficacy or control (Theme 3). In teaching, anxiety related to negative impact on others and role tension was often kept’ while in research anxiety related to negative impact on self was maintained from 2016 to 2017.

### GTAs pursuing academic versus non-academic careers differed in research anxieties

When asked in the second set of interviews if their career aspirations had changed, all GTAs said they had maintained the same primary career aspiration as the previous year. Biology GTAs who aspired to non-academic career aspirations often sought opportunities in government agencies or industry (n = 8) where research is the primary role. Some GTAs also indicated a desire to pursue teaching or outreach in museums, zoos, or K-12 classrooms (n = 4). Participants who espoused academic career pursuits included R1 professorships (n= 5), but mostly indicated positions in R2 or liberal arts colleges (n = 6), where teaching is a more prominent role.

Regardless of the career aspirations, negative impact on self and lack of self-efficacy still encompassed the most prevalent sources of anxiety in research (**Figure 5**). Similarly, anxiety related to negative impact on others in 2016 and role tension in 2017 were the most prevalent themes in teaching regardless of career aspirations. These observations, however, may be due to the small sample size representing different career aspiration groups. Despite these similarities, there appeared to be notable anxiety differences between career aspiration subgroups *within* research contexts. In Fall 2017, GTAs with non-academic career aspirations appeared to express higher percentages of anxiety related to role tensions and personal anxieties compared to GTAs with academic career aspirations (**Figure 5d**). GTAs who espoused these new anxieties in role tension or personal anxiety for 2017 described not having enough time to spend on research as they drew closer to graduation. For example, Eric articulated this anxiety related to pressure to spend all work hours on research, even when helping lab mates: “*pretty much everything that I’m doing I’m thinking I should be doing research all of the time*.”

**Figure 5:**
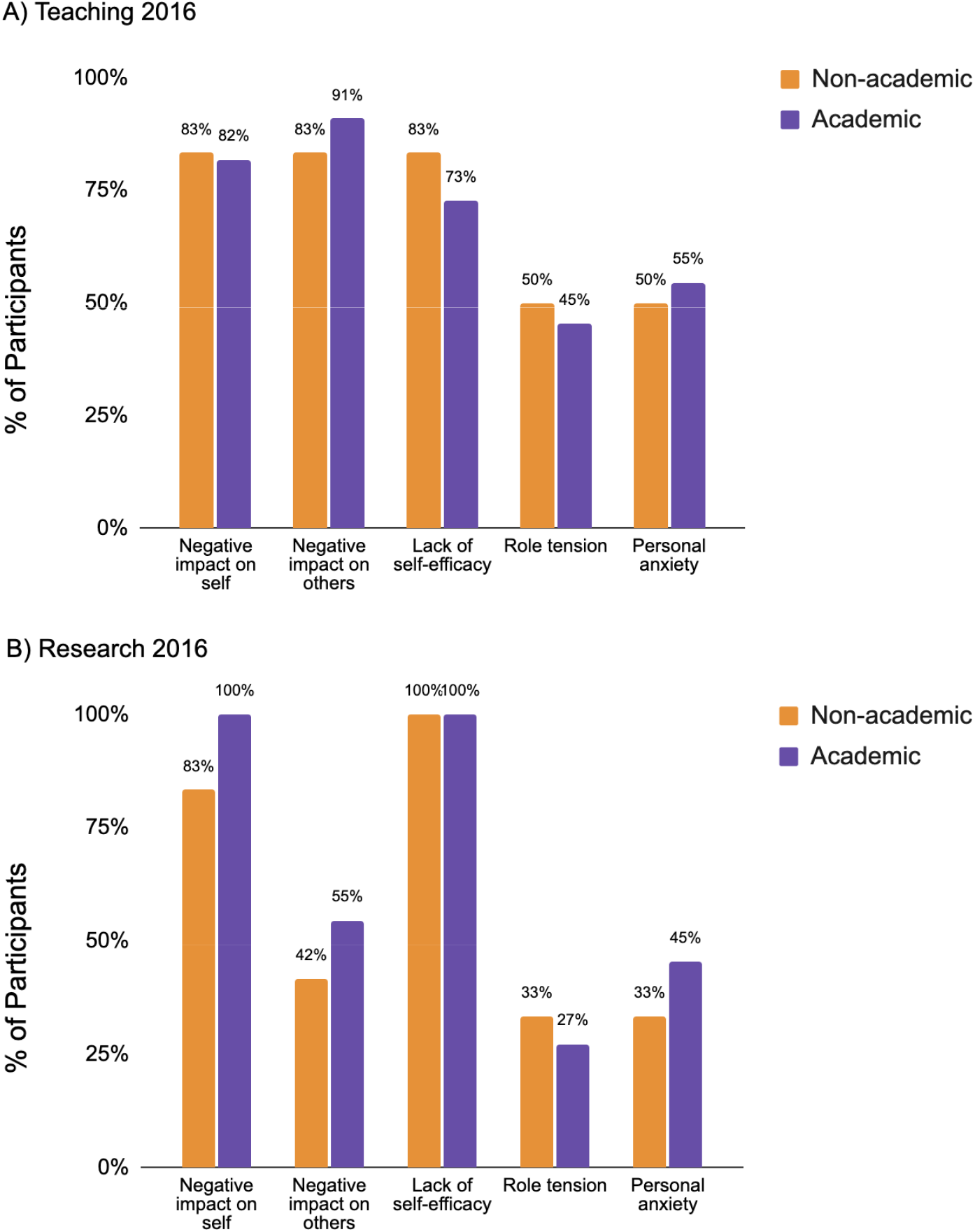
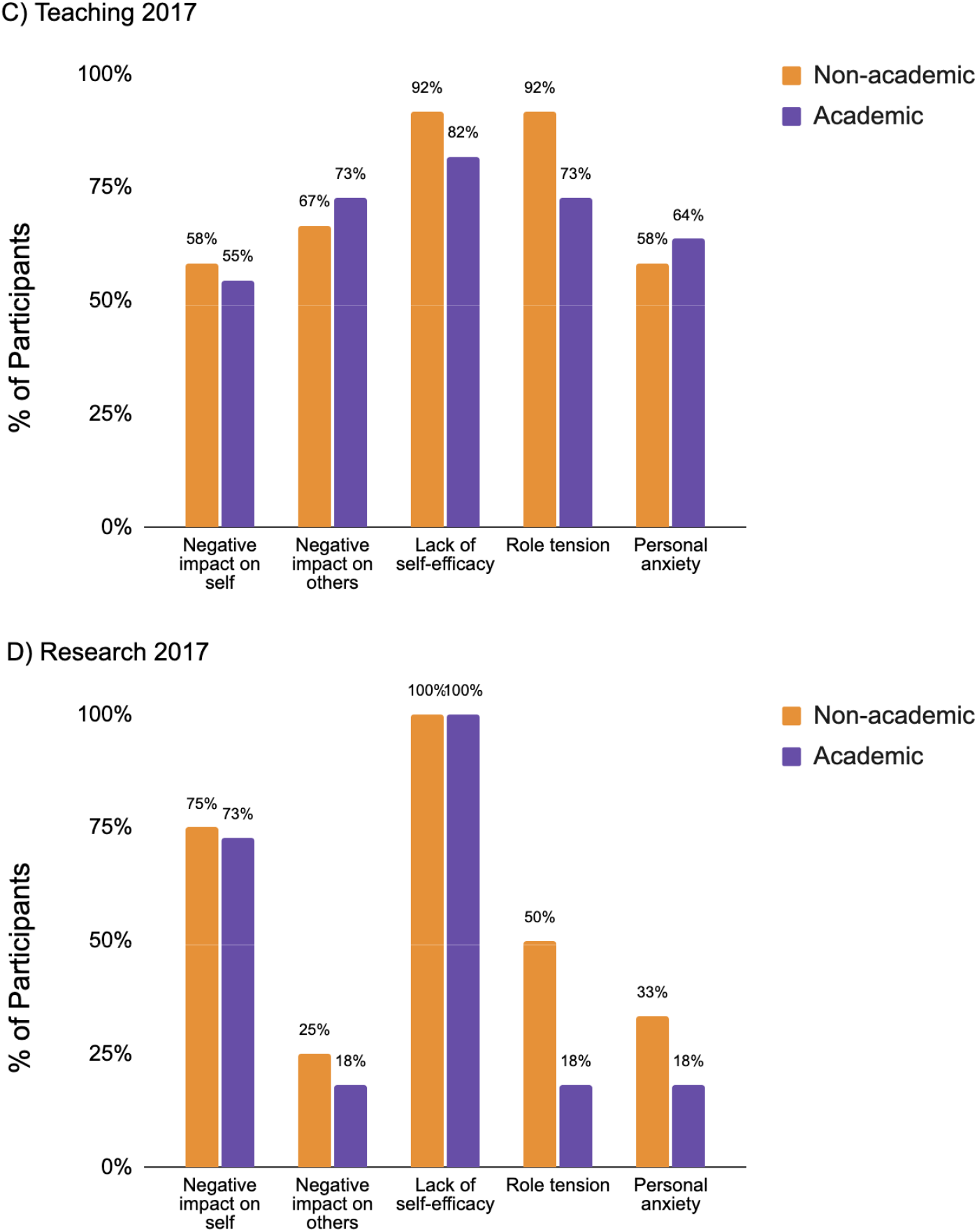
The percentage of participants (n=23) who exhibited each anxiety theme, comparing participants with non-academic (orange, n=12) or academic career interests (purple, n=11), for (**A**) teaching anxiety in Fall 2016, (**B**) research anxiety in Fall 2016, **(C)** teaching anxiety in Fall 2017, and **(D)** research anxiety in Fall 2017.

When comparing anxiety changes between GTAs with different career aspirations over time (**Figure 6**), trends differed based on teaching and research contexts. In teaching contexts, changes in anxieties over time were similar between GTAs with academic and non-academic career goals (**Figure 6a, c**). Anxieties related to negative impact on self, negative impact on others, lack of self-efficacy, and role tension changed similarly between academic and non-academic career aspiring GTAs. Changes in anxieties related to teaching varied in being kept, added, resolved, or never present, with anxiety related to lack of self-efficacy and negative impact on others often being most kept over time. Personal anxiety in teaching contexts had the only notable difference between subgroups, with non-academic GTAs having added more personal anxiety over time compared to academic GTAs. In research contexts, however, participants with academic career aspirations often resolved or never had their anxieties compared to those with non-academic aspirations (**Figure 6b, d**). Anxiety caused by lack of self-efficacy in research was kept among all GTAs with both academic and non-academic career aspirations over one year.

**Figure 6:**
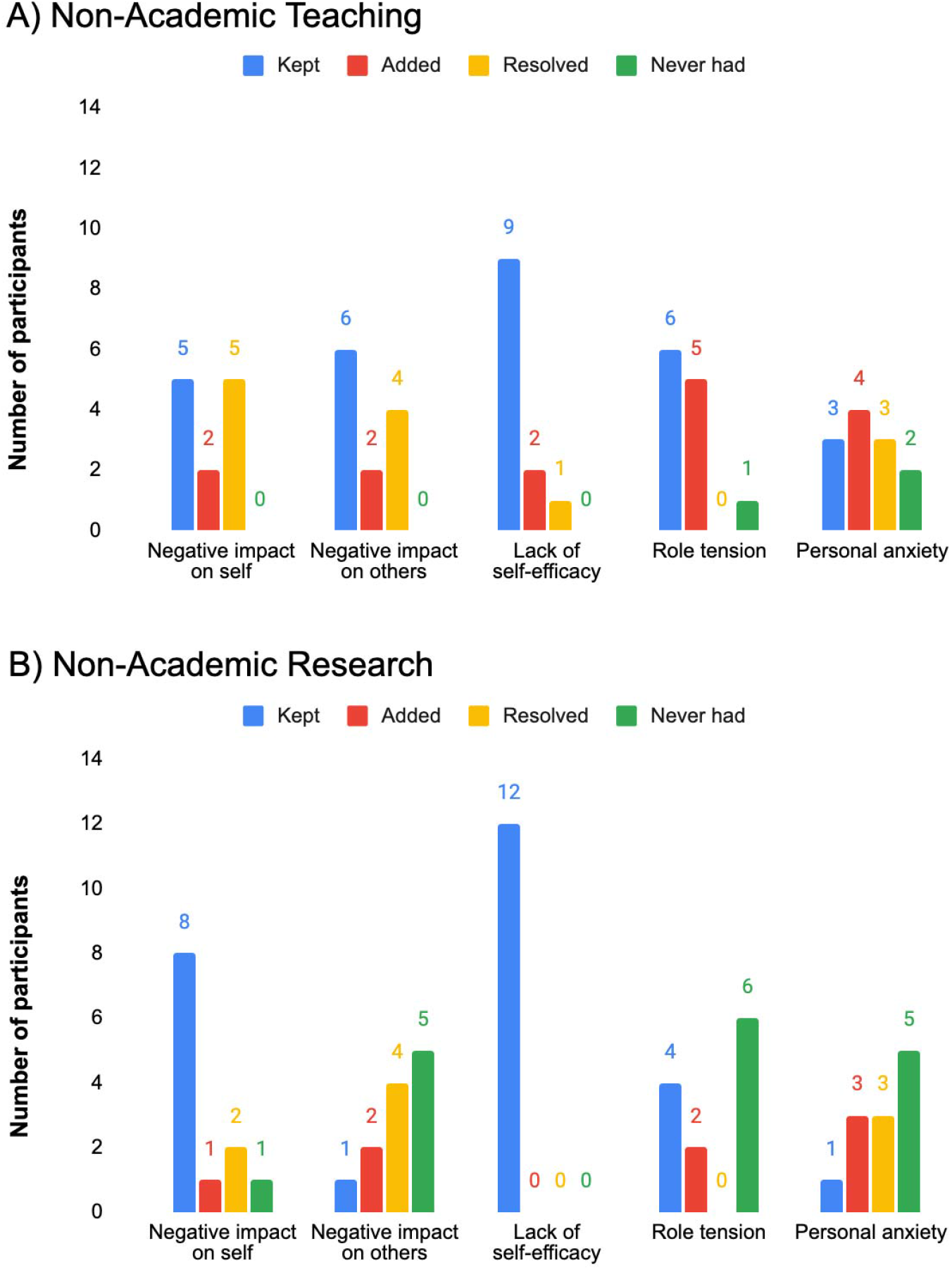
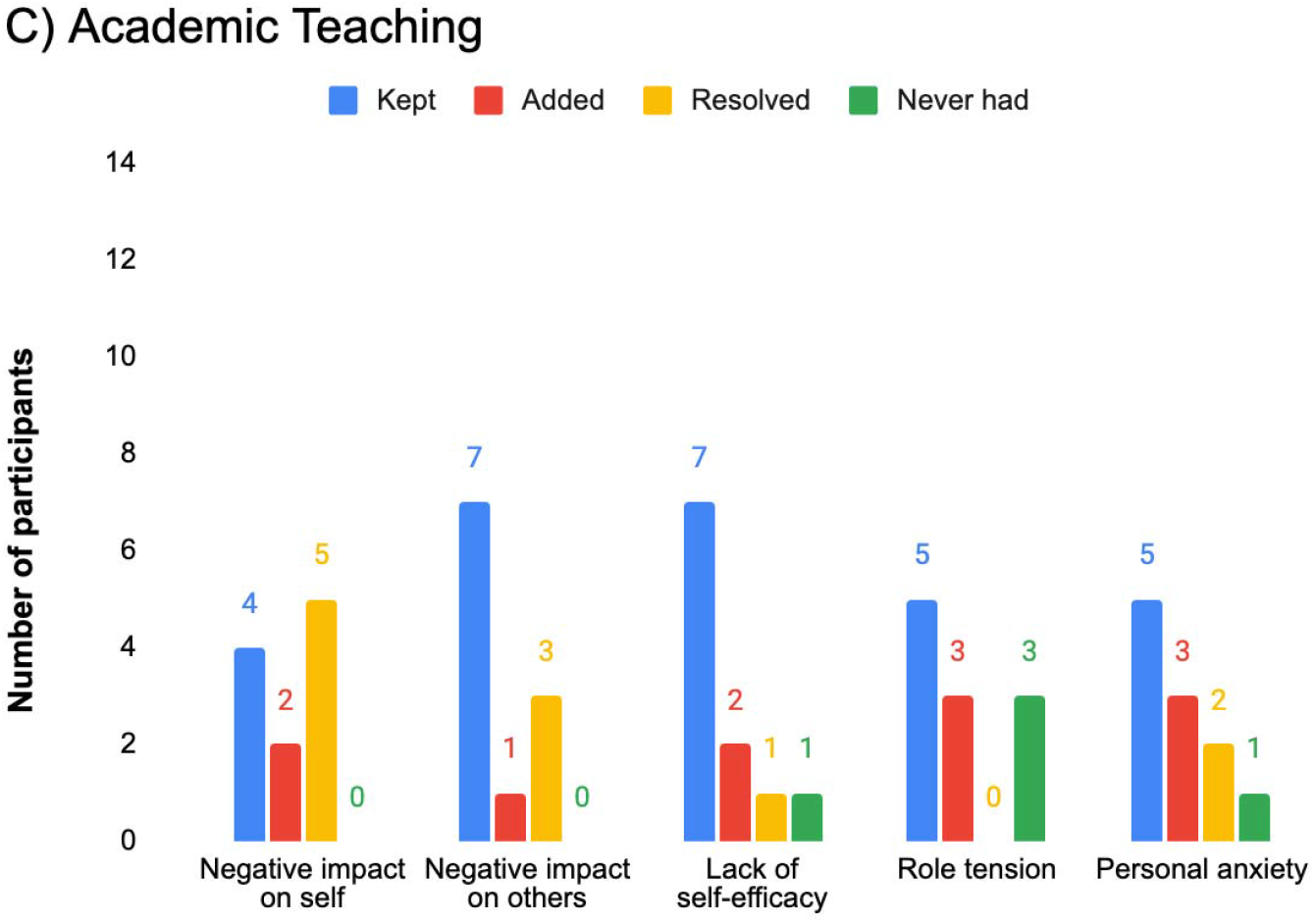
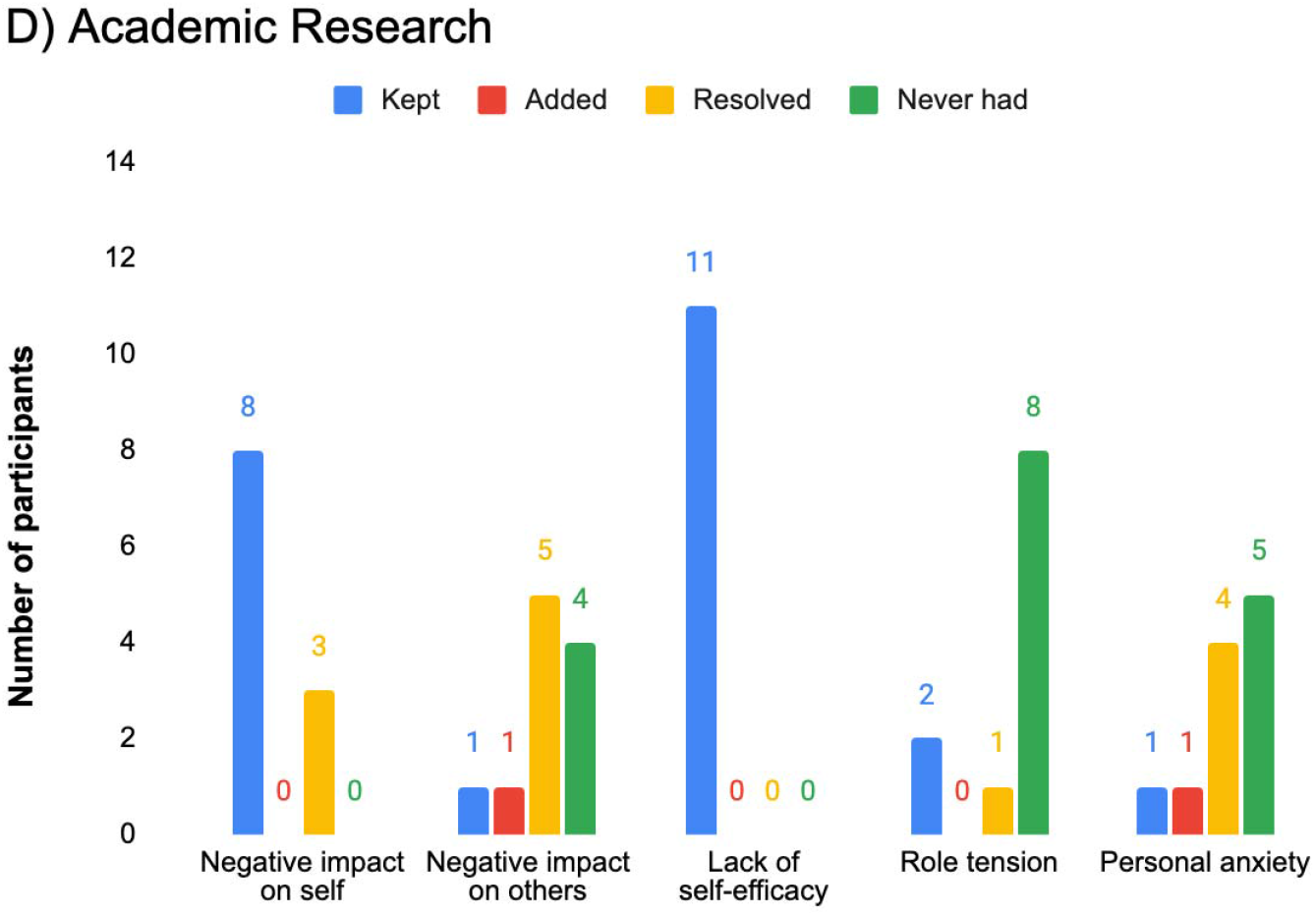
The number of Biology GTA participants which either kept, resolved, added, or never had that particular cause of anxiety from 2016 to 2017 for teaching and research between GTAs with non-academic (orange, n=12) and academic (purple, n=11) career.

## DISCUSSION

This study took a qualitative approach to probing what underlies anxieties specifically related to teaching and research roles for one group of Biology GTAs, how those sources of anxieties changed over time, and how the sources for teaching and anxiety compared based on career aspirations. The sources of anxieties which emerged among our Biology GTAs were related to cognitive and background variables on the SCCT model (**Figure 7**). Negative impact on self (Theme 1) and others (Theme 2), and role tension (Theme 4) are anxieties related to outcome expectations (e.g. “*what will happen to me if I fail to publish my research?*”). Self-efficacy (Theme 3) is a cognitive variable in SCCT. And lastly, personal anxieties (Theme 5) are related to SCCT’s background environmental or contextual variables. We found that teaching anxieties were most often related to a GTA’s perceived impact on others (e.g. students), while research anxieties were related to lack of self-efficacy. Role tension for teaching was the anxiety most likely to increase over time, while lack of self-efficacy was uniformly high over time for research especially. Interestingly, GTAs with academic career aspirations seemed to have less anxieties—either resolved or never had—related to research over time. This indicates that discussions of graduate student anxiety may need to consider the different roles of GTAs separately, in combination, over time, and with career aspirations in mind to gain a more nuanced understanding of GTA well-being.

**Figure 7:**
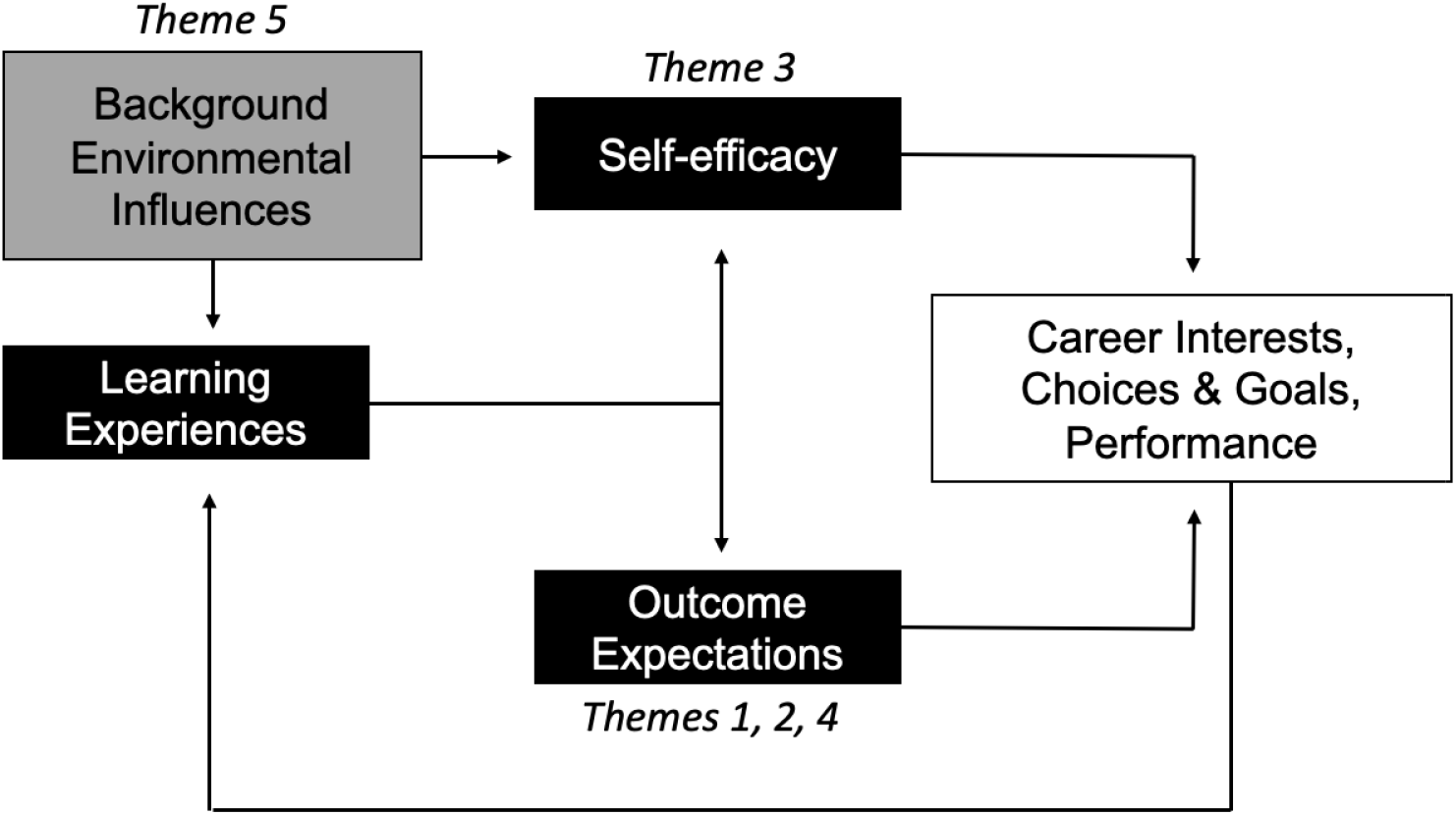
Modified Social Cognitive Career Theory (SCCT) model mapping the five anxiety themes found in this study with the cognitive and contextual factors which influence career interest development. Anxiety themes related to negative impact of self, negative impact on others, and role tension (Themes 1, 2, 4) were associated with outcome expectations, what GTAs expected to come out of certain professional interactions or decisions. Anxiety related to lack of self-efficacy (Theme 3) was related to a GTAs self-efficacy for a task and the perception that they could not complete it successfully or had no control to do so. Lastly, personal anxieties (Theme 5) are related a GTAs background influences such as departmental culture or personal life.

### Negative perception of others impacts GTA anxiety toward teaching and research

The perceptions of others, especially doctoral advisors and undergraduate students were related to two of the themes we identified in this study and is likely an important component of anxiety. Graduate school is not conducted in a void; students are constantly interacting with students, peers, and faculty and these interactions can influence how novices gradually adopt the identity of the profession (Adler and Adler 2005). It is no surprise that in these interactions are embedded fears of being covertly evaluated or compared (Adler and Adler 2005). Similar constructs have been found when studying anxiety related to active learning among science undergraduate students, with fear of negative evaluation a primary factor (Cooper, Downing, and Brownell 2018; Downing et al. 2020). Receiving positive feedback, recognition, and respect from faculty, peers, and students marks progress in graduate school socialization and likely reduces anxiety over time. Mentors, especially doctoral supervisors, play an integral role in successful socialization and sense of belonging to their mentees (McConnell, Geesa, and Lowery 2019). The student-advisor relationship is often the most important predictor of the satisfaction and persistence of a student’s doctoral experience (Devos et al. 2017; Hish et al. 2019; Hunter and Devine 2016; Zhao, Golde, and McCormick 2007), and influences the formation of a graduate student’s professional identity, the quality of their dissertation, professional network development, and available job prospects (Lovitts, 2001). Positive support and feedback from graduate student mentors can help bolster self-efficacy in teaching and research, helping them socialize into their professional identity. As Golde (2000) points out in studying graduate student experiences: “*pivotal in each story was the importance of a supportive advising relationship in helping students making progress toward their degree*.” (p. 219). In general, we found anxieties related to perceptions of others and its negative impact on self, decreased over time. It is possible that with successful socialization, GTAs may have less anxiety over how others perceive them. However, difficulties in socialization into the academic community would be expected to increase anxiety.

While the research anxiety related to perception of others was predominantly directed towards advisors, in a teaching context, this anxiety often related to undergraduates. GTAs focused on not letting down their students and establishing positive perceptions of themselves with their students. In many cases, GTAs may feel they need to work harder to establish their credibility in the classroom compared to faculty (Golish 1999; Hendrix 1995). In studies examining undergraduate perceptions of GTAs versus faculty, undergraduate students rated faculty members higher initially for being confident, enthusiastic, with more authority over the curriculum; while GTAs were rated higher for being nervous, uncertain, but having a more enjoyable instructional style (Kendall & Schussler, 2012; Kendall & Schussler, 2013). GTAs in this study often indicated a fear of improperly or accidentally conveying incorrect information to their students, thus negatively impacting their students’ learning. By providing teaching professional development opportunities, supportive preparatory meetings, or a system for structured teaching feedback, particularly for novice GTAs, teaching self-efficacy may increase, leading to reductions in anxieties related to perception of others (Connolly, Lee, and Savoy 2018; Reeves et al. 2018).

### Self-efficacy may be critical in reducing anxiety

Lack of self-efficacy (Bandura, 1988) was one of the most prominent anxieties among GTA participants for both teaching and research contexts, emerging often and consistently over time in this study. In research contexts, lack of self-efficacy was the only source of anxiety kept by all GTAs from 2016 to 2017. This comes as no surprise, as Bandura (1988) and others (Connolly, Lee, and Savoy 2018; DeChenne et al. 2015; Reeves et al. 2018) established that if self-efficacy for a task is high, then the anxiety toward said task is low. With forty percent of our Fall 2016 Biology GTA sample being first year GTAs, we may infer that self-efficacy may have been low for many of these students. Low self-efficacy occurs often with doing tasks such as teaching or doing research for the first time (Adler and Adler 2005). Several of the GTAs mentioned tasks being done for the first time in their interviews when asked to explain their anxieties.

Despite an expectation that self-efficacy may increase over time in graduate students, anxieties related to lack of self-efficacy for both teaching and research did not change over the year. In teaching settings, this can sometimes be related to a perceived lack of authority or autonomy of GTAs in the way a class or module is organized and taught (Muzaka 2009). Some GTAs also commented that they felt their voice in the organization, running of classes, and related issues was not important. Teaching assignments for graduate students are also not always stable. GTAs can be moved around to teach different courses, requiring new class or lab preparation. Research projects also change over time, requiring the learning of new skills at different stages of a project. In research, lack of self-efficacy over time may be attributed to part of their development as graduate students, where at different stages of the doctoral program, new tasks are encountered and self-efficacy must continually be built (Adler and Adler 2005; Gilmore et al. 2015). Austin (2002) indicates that factors such as the GTA’s locus of control (the extent to which a person perceives that they have the power to make decisions and manage the graduate experience) and the GTA’s sense of self-efficacy can affect the development of graduate students as prospective faculty. To build lasting teaching self-efficacy in GTAs, departments could make teaching assignments more stable over time, unless a GTA requests otherwise. To build research self-efficacy, advisors, departments, and institutions should consider how to better develop transferable skills in students, rather than having students fend for themselves (Hancock and Walsh 2016; Sinche et al. 2017). When newer educators and researchers enter into the program, they can be taught that expertise and self-efficacy grow with experience and professional development, which should be continually sought, in order to be less anxious about newly developing skills. Providing resources or programmatic interventions (Sinche et al. 2017) can go a long way to minimizing anxieties and supporting graduate students with a wide range of careers interests.

### Biology GTAs struggle with prioritizing research versus teaching resonsibilities and work-life balance

Tension between a GTA’s responsibilities in teaching and research are another point of anxiety, particularly related to time constraints. These tensions have been reported in previous work examining doctoral student’s perceptions of teaching, and in the development of GTA professional identity (Gilmore et al. 2015; McAlpine, Jazvac-Martek, and Hopwood 2009; Muzaka 2009). When Muzaka (2009) surveyed 10 GTAs and their perceptions of the most beneficial and problematic aspects of teaching, they found that the difficulties often related to time pressure. The majority of GTAs commented that teaching took considerable time away from their research and could delay timely graduation of their doctorate (Muzaka 2009). When McAlpine and colleagues (2009) studied Canadian Education doctoral students and their formation of academic identity, students reported that conducting research could pose difficulties in terms of the time available to do their work. Such role tension may be attributed to the messaging graduate students receive as to the most valued roles they are involved in by their advisors, department, and academic community interactions (Austin 2002; Austin and McDaniels 2006; Reid and Gardner 2020) e.g. research, then teaching, then service at some institutions. These findings are echoed in our Biology GTAs, which suggests that tensions between roles and responsibilities are to be expected within the experience and formation of professional identities in graduate school.

It should be noted, however, that investing time in pedagogy through teaching professional development (TPD) programs, may actually be beneficial for research preparation (Shortlidge and Eddy 2018) and does not delay graduate student time to graduate (Connolly et al. 2016). Gilmore and colleagues (2015) found that among the graduate students (n = 223) they surveyed, GTAs indicated a complimentary relationship between teaching and research (Gilmore et al. 2015). It is important for institutions to message to their students that teaching and teaching training can be incorporated into graduate programs without reducing students’ preparedness for a research career, and help students to see that these roles are not necessarily in conflict.

Lastly, personal anxiety (Theme 5) was expressed by many students in this study. According to a recent survey of over 6,000 graduate students, doctoral students hold great fears related to uncertainty of job prospects and difficulty maintaining a work–life balance (Woolston 2019). Hish et al. (2019) also found that personal anxieties (family and monetary stressors) correlated to depressive symptoms. Again, the process of graduate students’ socialization is not completed in isolation, nor can it be separated from a student’s personal life outside of the workplace. To encourage better school-work-life balance for students, doctoral advisors can model healthy role balances and flexibility in school and work schedules; while doctoral programs can continue to provide financial support, and support services tailored to specifically address doctoral student needs (Martinez et al. 2013; McConnell, Geesa, and Lowery 2019).

### Anxieties towards research declined over time for GTAs pursuing academic career aspirations

We found that individuals who wanted to pursue academic careers seemed to have resolved or never had anxiety pertaining to research by Fall 2017 compared to teaching. As expected from the SCCT model (Lent, Brown, and Hackett 1994), GTAs seemed to acclimate to the demands of academic research more easily if they were interested in a career in academia. As we found, with positive outcome expectations and learning experiences related to research, GTAs with academic career goals had overall fewer anxieties regarding research, despite lack of self-efficacy remaining a predominant factor related to anxiety. These GTAs also had some decreases in teaching anxieties. Over the span of their programs, STEM doctoral students often develop more interest in teaching undergraduate students, alongside their research interests, making teaching a predominant occupation (Connolly et al. 2016). The increasing interests in teaching may be driven by greater enjoyment of teaching or be a pragmatic shift because there are more teaching-focused positions available compared to research positions (Fuhrmann et al. 2011). Our GTAs who indicated a desire to pursue academia often included both a research and a teaching interest. Budding teaching interests could be related to decreases in research and teaching anxiety, as GTAs become more proficient in their research and teaching roles.

The changes in anxiety over time for GTAs with non-academic career aspirations reveal a more complex narrative, with no clear changes in anxieties over time. Surprisingly, the types of non-academic career aspirations listed by GTAs often included research in non-academic settings (e.g. government, industry, non-profit), suggesting that their anxieties in research could be specific to the academic research setting versus research in other contexts. Anxieties related to role tension increased among these GTAs, which can be expected if a GTAs is grappling with how to acquire the necessary skills to pursue their non-academic career interests which may not include teaching. Anxieties in teaching and research for GTAs with non-academic goals may be further exacerbated due to lack of departmental support in attaining these “non-traditional” career goals. O’Meara et al. (2014) found that departments can positively influence student agency by encouraging multiple career paths, providing information and financial support, and offering mentoring and guidance. By having advisors and departments support the formation and success of non-academic career paths for GTAs, anxieties for graduate students who want to pursue a non-academic career may be reduced.

### Improving GTA well-being: Building self-efficacy and coping strategies

A recent report by the National Academies of Sciences (2018) recommended that institutions and departments provide stronger support for graduate student mental health services. This call to action requires further in-depth research on the causes of graduate student mental health issues, so institutions and departments may make more informed decisions about *what* aspects of their programs may cause anxiety, *how* to support their graduate students at various points in their program and *which* graduate students need the greatest support. This work could inform these decisions and help identify groups (e.g. GTAs pursuing non-academic career paths) who may be more negatively impacted by anxiety and offer insight as to the most effective strategies of coping to alleviate destructive anxieties.

There are several ways graduate student well-being can be improved throughout their program. Through professional development opportunities, stress management techniques, teaching self-compassion, gratitude journaling, effective and supportive advisor mentoring, or encouraging counselling, academia can be a place for graduate students to maintain well-being (DeChenne, Enochs, and Needham 2012; Flinchbaugh et al. 2012; Golnaraghi 2016; National Academies of Sciences Engineering and Medicine 2018; Nicklin, Meachon, and McNall 2019; Reeves et al. 2018). For example, introducing coping strategies in new graduate student orientations or professional development may help to reduce anxieties from the outset. Byers et al. (2014) found five major themes emerge during their interviews on survival strategies for doctoral students: compartmentalization of life, outside support systems, justification for participation in program, emotional status, and structure of program (Byers et al. 2014). By emphasizing to graduate students, especially incoming novice graduate students from both masters or doctoral programs, that these aspects may help one maintain mental health, we can improve the graduate student experience and reduce attrition of vulnerable groups from academia.

### Limitations: Not all anxiety is equal

Similar to previous studies in this research group (Chen and Schussler, in prep), this research may lead the reader to assume that anxiety is not desirable, and negatively impacts an individual’s well-being and daily life functioning. However, we acknowledge that not all anxiety has the same effect, and that some anxiety can actually be a positive force for productivity and creativity. Yerkes and Dodson (1908) established that there is an ideal threshold in which “arousal” or anxiety can actually increase productivity (Cohen 2011; Pelton 2014). Too much and too little anxiety are both likely impediments to achievement, and to a certain extent, some level of doubt or lack of confidence may provide significant impetus to improving effectiveness such as in teaching (Wheatley 2005). Though we recognize some anxiety as motivating, anxiety can only help individual well-being if positive coping strategies are present. Therefore, for the future, to distinguish between negative and positive anxieties, we hope to additionally examine how individuals cope with each anxiety. If the coping is constructive and effective, anxiety may actually be productive. If the coping involves avoidance and is destructive, that anxiety may have a negative impact. By examining these anxieties in a longitudinal framework, we can also investigate whether coping and anxiety becomes more or less productive over time.

### Conclusions

To tackle the graduate student anxiety epidemic, there must be a better understanding of what makes students anxious in order to propose methods to reduce that anxiety. This study provided an important first step to help understand how others in the academic context, self-efficacy, and personal perceptions play a role in mental health, and how these may differ with the varying teaching and research roles graduate students play at institutions. In some cases, the anxieties students expressed such as role tension or work-life balance may be symptomatic of systemic structural problems in academia that will be hard to change. However, professional development activities or training opportunities for GTAs pose a tangible, effective method for institutions and departments to consider mitigating anxiety. Professional development workshops may also provide efficacious coping strategies to regulate external stressors which cause anxiety and encourage greater sense of community. In this way, universities can start to take responsibility for not only training graduate students in the academic discipline, but also helping them navigate the cognitive and emotional outcomes that are a common part of seeking a higher degree.

## ACKNOWLEDGEMENTS

Thank you to Caroline Wienhold, Maryrose Weatherton, Randall Small, Sarah Andrews, Nicole Chodkowski, Robert Furrow, and Brie Tripp for their comments on earlier versions of the manuscript. We appreciate the feedback from two anonymous reviewers in subsequent versions of the manuscript. Thank you to the Ecology and Evolutionary Department for providing research funds to support our incentives. Lastly, thank you to all GTA participants for taking part in this research.

## REFERENCES

Adler, Patricia A., and Peter Adler. 2005. “The Identity Career of the Graduate Student: Professional Socialization to Academic Sociology.” American Sociologist 36(2): 11–27.

American Psychological Association. 2020. “Definition of Anxiety.”: Psychological Topics. https://www.apa.org/topics/anxiety/ (March 10, 2020).

Austin, Ann E. 2002. “Preparing the Next Generation of Faculty.” The Journal of Higher Education 73(1): 94–122.

Austin, Ann E., and Melissa McDaniels. 2006. “Preparing the Professoriate of the Future: Graduate Student Socialization for Faculty Roles.” In Higher Education: Handbook of Theory and Research,.

Bandura, Albert. 1988. “Self-Efficacy Conception of Anxiety.” Anxiety Research 7(1): 77–98.

Bandura, Albert.. 1993. “Perceived Self-Efficacy in Cognitive Development and Functioning.” Educational Psychologist 28(2): 117–48. http://www.tandfonline.com/doi/abs/10.1207/s15326985ep2802_3.

Barry, K. M. et al. 2018. “Psychological Health of Doctoral Candidates, Study-Related Challenges and Perceived Performance.” Higher Education Research and Development 37(3): 468–83.

Blake, C. E. et al. 2007. “Classifying Foods in Contexts: How Adults Categorize Foods for Different Eating Settings.” Appetite 49(2): 500–510.

Boyatzis, Richard. 1998. Thematic analysis and code development *Transforming Qualitative Information*.

Braun, Virginia, and Victoria Clarke. 2006. “Using Thematic Analysis in Psychology.” Qualitative Research in Psychology.

Brinkman, Stacy N., and Arianne A. Hartsell-Gundy. 2012. “Building Trust to Relieve Graduate Student Research Anxiety.” Public Services Quarterly 8(1): 26–39.

Bucher, Rue, and Joan G Stelling. 1977. Becoming professional. Becoming Professional. Oxford, England: Sage.

Byers, Valerie Tharp et al. 2014. “Survival Strategies: Doctoral Students’ Perceptions of Challenges and Coping Methods.” International Journal of Doctoral Studies 9: 109–36.

Charmaz, Kathy. 2006. International Journal of Qualitative Studies on Health and Wellbeing *Constructing Grounded Theory: A Practical Guide through Qualitative Analysis (Introducing Qualitative Methods Series)*.

Cho, YoonJung, Myoungsook Kim, Marilla D. Svinicki, and Mark Lowry Decker. 2011. “Exploring Teaching Concerns and Characteristics of Graduate Teaching Assistants.” Teaching in Higher Education 16(3): 267–79. http://www.tandfonline.com/doi/abs/10.1080/13562517.2010.524920.

Clair, Rebekah St et al. 2017. “The New Normal: Adapting Doctoral Trainee Career Preparation for Broad Career Paths in Science.” PLoS ONE 12(5).

Coates, Thomas J., and Carl E. Thoresen. 1976. “Teacher Anxiety: A Review with Recommendations.” Review of Educational Research 46(2): 159–84.

Cohen, Ronald A. 2011. “Yerkes–Dodson Law.” In Encyclopedia of Clinical Neuropsychology,.

Connolly, Mark R., You Geon Lee, and Julia N. Savoy. 2018. “The Effects of Doctoral Teaching Development on Early-Career STEM Scholars’ College Teaching Self-Efficacy.” CBE Life Sciences Education 17(1): 1–15.

Connolly, Mark R, Julia N Savoy, You-geon Lee, and Lucas B Hill. 2016. Building a Better Future STEM Faculty: How Teaching Development Programs Can Improve Undergraduate Education. Madison, WI.

Conrad, Lettie Y., and Virginia M. Tucker. 2019. “Making It Tangible: Hybrid Card Sorting within Qualitative Interviews.” Journal of Documentation 75(2): 397–416.

Cooper, Katelyn M., Virginia R. Downing, and Sara E. Brownell. 2018. “The Influence of Active Learning Practices on Student Anxiety in Large-Enrollment College Science Classrooms.” International Journal of STEM Education 5(1).

DeChenne, Sue Ellen, Larry G. Enochs, and Mark Needham. 2012. “Science, Technology, Engineering, and Mathematics Graduate Teaching Assistants Teaching Self-Efficacy.” Journal of the Scholarship of Teaching and Learning 12(4): 102–23. http://josotl.indiana.edu/article/view/2131.

DeChenne, Sue Ellen, Natalie Koziol, Mark Needham, and Larry Enochs. 2015. “Modeling Sources of Teaching Self-Efficacy for Science, Technology, Engineering, and Mathematics Graduate Teaching Assistants.” CBE Life Sciences Education 14(3): 1–14.

Denise Kendall, K., and Elisabeth E. Schussler. 2012. “Does Instructor Type Matter? Undergraduate Student Perception of Graduate Teaching Assistants and Professors.” CBE Life Sciences Education 11(2): 187–99.

Devos, Christelle et al. 2017. “Doctoral Students’ Experiences Leading to Completion or Attrition: A Matter of Sense, Progress and Distress.” European Journal of Psychology of Education 32(1): 61–77.

Downing, Virginia R. et al. 2020. “Fear of Negative Evaluation and Student Anxiety in Community College Active-Learning Science Courses.” CBE Life Sciences Education 19(2): 1–16.

El-Ghoroury, Nabil Hassan, Daniel I. Galper, Abere Sawaqdeh, and Lynn F. Bufka. 2012. “Stress, Coping, and Barriers to Wellness among Psychology Graduate Students.” Training and Education in Professional Psychology 6(2): 122–34.

Evans, Teresa M et al. 2018. “Evidence for a Mental Health Crisis in Graduate Education.” Nature Biotechnology 36(3): 282–84. http://dx.doi.org/10.1038/nbt.4089.

Flinchbaugh, Carol L., E. Whitney G. Moore, Young K. Chang, and Douglas R. May. 2012. “Student Well-Being Interventions: The Effects of Stress Management Techniques and Gratitude Journaling in the Management Education Classroom.” Journal of Management Education 36(2): 191–219.

Fuhrmann, C N, D G Halme, P. S. O’Sullivan, and B Lindstaedt. 2011. “Improving Graduate Education to Support a Branching Career Pipeline: Recommendations Based on a Survey of Doctoral Students in the Basic Biomedical Sciences.” CBE Life Sciences Education 10(3): 239–49.

Gardner, Grant, and M. Gail Jones. 2011. “Pedagogical Preparation of the Science Gaduate Teaching Assistant: Challenges and Implications.” Science Educator 20(2): 31–41.

Gilmore, Joanna et al. 2015. “Feeding Two Birds with One Scone? The Relationship between Teaching and Research for Graduate Students across the Disciplines.” International Journal of Teaching and Learning in Higher Education 27(1): 25–41. http://www.isetl.org/ijtlhe/.

Glaser, B., and A. Strauss. 1967. “The Discovery of Grounded Theory. 1967.” Weidenfield & Nicolson, London. https://www.coursera.org/.

Golde, Chris M. 2000. “Should i Stay or Should i Go? Student Descriptions of the Doctoral Attrition Process.” Review of Higher Education.

Golde, Chris M.. 2005. “The Role of the Department and Discipline in Doctoral Student Attrition: Lessons from Four Departments.” The Journal of Higher Education 76(6): 669–700. http://muse.jhu.edu/content/crossref/journals/journal_of_higher_education/v076/76.6golde.html.

Golish, Tamara D. 1999. “Students’ Use of Compliance Gaining Strategies with Graduate Teaching Assistants: Examining the Other End of the Power Spectrum.” Communication Quarterly 47(1): 12–32.

Golnaraghi, Golnaz. 2016. “The Power of Self-Compassion in the Doctoral Journey.”: 129–41.

Gregory, Donald G., and Barbara E. Lovitts. 2003. “Leaving the Ivory Tower: The Causes and Consequences of Departure from Doctoral Study.” Contemporary Sociology.

Hancock, Sally, and Elaine Walsh. 2016. “Beyond Knowledge and Skills: Rethinking the Development of Professional Identity during the STEM Doctorate.” Studies in Higher Education 41(1): 37–50.

Hendrix, Katherine Grace. 1995. “Preparing Graduate Teaching Assistants (GTAs) to Effectively Teach the Basic Course.” Annual Meeting of the Southern States Communication Association.

Hish, Alexander J. et al. 2019. “Applying the Stress Process Model to Stress–Burnout and Stress–Depression Relationships in Biomedical Doctoral Students: A Cross-Sectional Pilot Study.” CBELife Sciences Education 18(4): 1–11.

Hunter, Karen H., and Kay Devine. 2016. “Doctoral Students’ Emotional Exhaustion and Intentions to Leave Academia.” International Journal of Doctoral Studies 11: 35–61.

Kajfez, Rachel L., and Lisa D McNair. 2014. “Graduate Student Identity: A Balancing Act between Roles.” In 121st ASEE Annual Conference & Exposition, ed. American Society for Engineering Education. Indianapolis, IN, 1–16.

Kendall, K. Denise, and Elisabeth E. Schussler. 2013. “Evolving Impressions: Undergraduate Perceptions of Graduate Teaching Assistants and Faculty Members over a Semester.” CBE Life Sciences Education 12(1): 92–105.

Lane, A. K., J. Skvoretz, et al. 2019. “Investigating How Faculty Social Networks and Peer Influence Relate to Knowledge and Use of Evidence-Based Teaching Practices.” International Journal of STEM Education 6(1).

Lane, A.K., Carlton Hardison, Ariana Simon, and Tessa C Andrews. 2019. “A Model of the Factors Influencing Teaching Identity among Life Sciences Doctoral Students.” Journal of Research in Science Teaching 56(2): 141–62. http://doi.wiley.com/10.1002/tea.21473.

Larson, Richard C., Navid Ghaffarzadegan, and Yi Xue. 2014. “Too Many PhD Graduates or Too Few Academic Job Openings: The Basic Reproductive Number R0 in Academia.” Systems Research and Behavioral Science 31(6): 745–50.

LeBreton, James M., and Jenell L. Senter. 2008. “Answers to 20 Questions about Interrater Reliability and Interrater Agreement.” Organizational Research Methods 11(4): 815–52.

Lent, Robert W., Steven D. Brown, and Gail Hackett. 1994. “Toward a Unifying Social Cognitive Theory of Career and Academic Interest, Choice, and Performance.Pdf.”

Lent, Robert W, Steven D Brown, and Gail Hackett. 2000. “Contextual Supports and Barriers to Career Choice: A Social Cognitive Analysis.” Journal of Counseling Psychology 47(1): 36–49. http://doi.apa.org/getdoi.cfm?doi=10.1037/0022-0167.47.1.36.

Levecque, Katia et al. 2017. “Work Organization and Mental Health Problems in PhD Students.” Research Policy 46(4): 868–79. http://dx.doi.org/10.1016/j.respol.2017.02.008.

Lindholm, Jennifer A. 2004. “Pathways to the Professoriate: The Role of Self, Others, and Environment in Shaping Academic Career Aspirations.” Journal of Higher Education 75(6): 603–35.

Lovitts, Barbara E. 2001. “Leaving the Ivory Tower: The Causes and Consequences of Departure from Doctoral Study.” Causes and consequences of departure from doctoral study.

Martinez, Edna;, Chinasa; Ordu, Matthew R.; Della Sala, and Adam McFarlane. 2013. “Striving to Obtain a School-Work-Life-Balance: The Full-Time Doctoral Student.” International Journal of Doctoral Studies 8: 39–59.

McAlpine, Lynn, Marian Jazvac-Martek, and Nick Hopwood. 2009. “Doctoral Student Experience in Education: Activities and Difficulties Influencing Identity Development.” International Journal for Researcher Development 1(1): 97–109.

McConnell, Kat, Rachel Louise Geesa, and Kendra Lowery. 2019. “Self-Reflective Mentoring: Perspectives of Peer Mentors in an Education Doctoral Program.” International Journal of Mentoring and Coaching in Education 8(2): 86–101.

McHugh, Marry L. 2012. “Interrater Reliability: The Kappa Statistic.” Biochemia Medica: 276–82. http://www.biochemia-medica.com/node/501.

Merriam, Sharan B. 1995. “What Can You Tell from an N of 1? Issues of Validity and Reliability in Qualitative Research.” PAACE Journal of Lifelong Learning 4: 51–60.

Merriam, Sharan B, and Elizabeth J. Tisdell. 2016. Qualitative Research: A Guide to Design and Implementation. 4th ed. ed. Elizabeth J Tisdell. Newark: John Wiley & Sons, Incorporated.

Mousavi, Maral P S et al. 2018. “Stress and Mental Health in Graduate School: How Student Empowerment Creates Lasting Change.”

Muzaka, Valbona. 2009. “The Niche of Graduate Teaching Assistants (GTAs): Perceptions and Reflections.” Teaching in Higher Education 14(1): 1–12.

Nagy, Gabriela A. et al. 2019. “Burnout and Mental Health Problems in Biomedical Doctoral Students.” CBE life sciences education 18(2): ar27.

National Academies of Sciences Engineering and Medicine. 2018. Graduate STEM Education for the 21st Century Graduate STEM Education for the 21st Century. eds. Alan Leshner and Layne Scherer. Washington, D.C.: National Academies Press. https://www.nap.edu/catalog/25038.

Nicklin, Jessica M, Emily J Meachon, and Laurel A. McNall. 2019. “Balancing Work, School, and Personal Life among Graduate Students: A Positive Psychology Approach.” Applied Research in Quality of Life 14(5): 1265–86.

OECD. 2014. “Who Are the Doctorate Holders and Where Do Their Qualifications Lead Them?” Education Indicators in Focus.

Parsons, Jane S. 1973. “Assessment of Anxiety About Teaching Using the Teaching Anxiety Scale: Manual and Research.” Research and Development Center for Teacher Education: 1–55.

Pekrun, Reinhard, Anne C. Frenzel, Thomas Goetz, and Raymond P. Perry. 2007. “The Control-Value Theory of Achievement Emotions. An Integrative Approach to Emotions in Education.” In Emotion in Education, Educational Psychology, 13–36.

Pelton, Julie A. 2014. “Assessing Graduate Teacher Training Programs: Can a Teaching Seminar Reduce Anxiety and Increase Confidence?” Teaching Sociology 42(1): 40–49. http://tso.sagepub.com/cgi/doi/10.1177/0092055X13500029.

Prieto, Loreto R., and Elizabeth M. Altmaier. 1994. “The Relationship of Prior Training and Previous Teaching Experience to Self-Efficacy among Graduate Teaching Assistants.” Research in Higher Education 35(4): 481–97.

Prieto, Loreto R., and Karen R. Scheel. 2008. “Teaching Assistant Training in Counseling Psychology.” Counselling Psychology Quarterly 21(1): 49–59.

Reeves, Todd D. et al. 2018. “Does Context Matter? Convergent and Divergent Findings in the Cross-Institutional Evaluation of Graduate Teaching Assistant Professional Development Programs.” CBE Life Sciences Education 17(1): 1–13.

Reeves, Todd D et al. 2016. “A Conceptual Framework for Graduate Teaching Assistant Professional Development Evaluation and Research.” 15: 1–9.

Reid, Joshua W., and Grant E. Gardner. 2020. “Navigating Tensions of Research and Teaching: Biology Graduate Students’ Perceptions of the Research–Teaching Nexus within Ecological Contexts.” CBE Life Sciences Education 19(3): 1–16.

Roach, K. David. 2003. “Teaching Assistant Anxiety and Coping Strategies in the Classroom.” Communication Research Reports 20(June): 81–89.

Saldaña, Johnny. 2012. “An Introduction to Codes and Coding.” The Coding Manual for Qualitative Researchers (2006): 1–8.

Schwandt, Thomas. 2007. The SAGE Dictionary of Qualitative Inquiry *Thematic Analysis*. http://srmo.sagepub.com/view/the-sage-dictionary-of-qualitative-inquiry/SAGE.xml.

Shortlidge, Erin E., and Sarah L. Eddy. 2018. “The Trade-off between Graduate Student Research and Teaching: A Myth?” PLoS ONE 13(6): 4–6.

Sinche, Melanie et al. 2017. “An Evidence-Based Evaluation of Transferrable Skills and Job Satisfaction for Science PhDs.” PloS one 12(9): e0185023.

Spradley, James P. 1979. “The Ethnographic Interview.”

Strauss, Anselm, and Juliet Corbin. 2008. 3 Basics of Qualitative Research Grounded Theory Procedures and Techniques *Basics of Qualitative Research: Techniques and Procedures for Developing Grounded Theory*. http://srmo.sagepub.com/view/basics-of-qualitative-research/SAGE.xml.

Sundberg, Marshall D, Joseph E Armstrong, and E William Wischusen. 2005. “A Reappraisal of the Status of Introductory Biology Laboratory Education in U.S. Colleges & Universities.” American Biology Teacher 67(9): 525–29. http://prx.library.gatech.edu/login?url=http://search.ebscohost.com/login.aspx?direct=true&db=eric&AN=EJ796369&site=ehost-live%5Cnhttp://www.nabt.org.

Weller, Susan C, and A Kimball Romney. 1988. Systematic Data Collection. Los Angeles: SAGE Publications Inc.

Wenger, Etienne. 1998. “Communities of Practice: Learning, Meaning, and Identity.” Systems thinker.

Wheatley, Karl F. 2005. “The Case for Reconceptualizing Teacher Efficacy Research.” Teaching and Teacher Education 21(7): 747–66.

Winstone, Naomi, and Darren Moore. 2017. “Sometimes Fish, Sometimes Fowl? Liminality, Identity Work and Identity Malleability in Graduate Teaching Assistants.” Innovations in Education and Teaching International 54(5): 494–502. http://dx.doi.org/10.1080/14703297.2016.1194769.

Woolston, Chris. 2019. “PhD Poll Reveals Fear and Joy, Contentment and Anguish.” Nature 575(13 November 2019): 403–6. https://www.nature.com/articles/d41586-019-03459-7.

Zhao, Chun Mei, Chris M. Golde, and Alexander C. McCormick. 2007. “More than a Signature: How Advisor Choice and Advisor Behaviour Affect Doctoral Student Satisfaction.” Journal of Further and Higher Education.

