## Supplemental Materials for "Finding a Balance: Characterizing Teaching and Research Anxieties in Biology Graduate Teaching Assistants (GTAs)"

**SUPPLEMENTAL MATERIAL**

**INTERVIEW PROTOCOL**

*Before you read this, read the informed consent for interview participants, and have them sign the general consent and audio record consent on the back. Then, proceed to this sheet.*

The questions I am going to ask you today are about four specific topics regarding your teaching: your experience in teaching, perceptions of teaching and research anxiety, the coping strategies you enact (this just means to manage your anxiety level to decrease it), and your current career aspirations. There are no right or wrong answers to these questions, and these responses are completely confidential – I just want to gain your perspective about these ideas!

**A) Career/Identity**

The first thing I’d like you to do is to take a moment and write down your top two current career aspirations. We’ll revisit this again at the end.

**B) Teaching experience, knowledge, and attitudes**

This portion of the interview will cover general experience and perceptions of teaching.

******** *Interviewees will have a scale in front of them with three pieces of post-it paper to rank: Experience, Knowledge and Attitude. They will be ranking themselves and explaining their choices.*

1. Do you consider yourself to be an experienced teacher? On a scale 1 to 10, 1 indicating little experience and 10 being highly experienced, how would you rate yourself?
2. Based on your experience, on the same scale, 1 representing little knowledge and 10 being highly knowledgeable, how would you rate yourself on knowledge in teaching. This can be from pedagogical knowledge, to assessment design, etc.
3. On a scale 1 to 10, 1 indicating very negative and 10 being very positive, how would you rate your attitude toward teaching.
4. Please explain your choices.

**C) Teaching and Research Anxiety**

5. Before jumping into the next question, I’d like you to take a minute and list a few things that make you anxious (if you have any) about teaching and research.

***have them complete this on the prepared piece of paper with the career aspirations as well*

Thank you. To continue, we’re going to look at these cards and identify aspects of teaching that may cause you anxiety. This protocol is still in development, so feel free to create your own cards and add to the list.

***Layout cards in front of participants randomly. Use cards to have them select things that make them anxious (or create own cards) and have them rank (1 being most anxious).* ***If they do not pick a card, skip to 5d.***

a. What about each choice specifically that make you anxious?
b. How do you think anxiety impacts your teaching for each? +/-/0
c. How you cope (if you do cope) with these things for each?
d. You did not choose any cards that make you anxious, why is that? Do you consider yourself not a very anxious person or have specific coping strategies you use?
6. Does the research you are conducting as part of your graduate program ever make you feel anxious? And for this question, we’re going to look at these cards and identify aspects that may cause you anxiety. Not all cards will represent what you are anxious about, so again, feel free to add to the list.

***Layout card in front of participants randomly. Use cards to have them select things that make them anxious (or create own cards) and have them rank (1 being most anxious).* ***If they do not pick a card, skip to 6d.***

a. Generally, what about these choices specifically make you anxious?
b. How do you think anxiety impacts your research? +/-/0
c. How do you cope (if you do) with these things?
d. You did not choose any cards that make you anxious, why is that? Do you consider yourself not a very anxious person or have specific coping strategies you use?

*****If there are rankings for BOTH teaching and research anxiety proceed to Q7. If not continue to Q8.***

7. We have this list of things that make you anxious about teaching, and this list of things that make you anxious about research. What I want you to do now is rank ALL these cards according to what causes you the most down to the least anxiety OVERALL in grad school.

***use cards to have them ranks things that make them anxious (or create own cards) from the first two sets.*

a. So it looks like [teaching / research / a mixture of both] cause you the most anxiety in grad school. Why is that?

b. Does your teaching anxiety impact your research anxiety? If so, how?

c. Does your research anxiety impact your teaching anxiety? If so, how?

d. Do you have any other sources of anxiety in grad school?

8. Would you want graduate school (teaching and research) to cause you no anxiety? Why or why not?

**D) Revisiting career aspirations/identity and attrition**

9. Revisiting your top career aspirations, what are they and why did you choose them? Assume the world is your oyster—would they be the same? i.e. not influences by anything like the job market, etc.
10. Does your anxiety about teaching/research/both make you second guess or question your career options? And if so why?
11. How likely could the anxiety that you feel in teaching, research, or both cause you to consider leaving the program?
12. Do you have any other thoughts about being a graduate student, anxiety, teaching, research, etc. you would like to share?

Thank you!

*Have students complete the compensation form and then sign the compensation consent line on the informed consent sheet.*

**A) Career aspirations**

1.

2.

**C) Anxieties**

| **Teaching Anxiety** | **Research Anxiety** |
| --- | --- |

**Cards for Teaching Anxiety (Blue Cards)**

Student issues/Emergency situations

| Anxiety_T | Anxiety impact on teaching (+/-/0)? | Anxiety_G |
| --- | --- | --- |

Time away from other priorities

| Anxiety_T | Anxiety impact on teaching (+/-/0)? | Anxiety_G |
| --- | --- | --- |

Supporting effective student learning

| Anxiety_T | Anxiety impact on teaching (+/-/0)? | Anxiety_G |
| --- | --- | --- |

Feedback/complaints about teaching

| Anxiety_T | Anxiety impact on teaching (+/-/0)? | Anxiety_G |
| --- | --- | --- |

Interactions with teaching supervisors

| Anxiety_T | Anxiety impact on teaching (+/-/0)? | Anxiety_G |
| --- | --- | --- |

| Anxiety_T | Anxiety impact on teaching (+/-/0)? | Anxiety_G |
| --- | --- | --- |

Being observed teaching by a peer

| Anxiety_T | Anxiety impact on teaching (+/-/0)? | Anxiety_G |
| --- | --- | --- |

Living off my stipend

| Anxiety_T | Anxiety impact on teaching (+/-/0)? | Anxiety_G |
| --- | --- | --- |

Reading student evaluations

| Anxiety_T | Anxiety impact on teaching (+/-/0)? | Anxiety_G |
| --- | --- | --- |

Attending TA Meetings

| Anxiety_T | Anxiety impact on teaching (+/-/0)? | Anxiety_G |
| --- | --- | --- |

Being observed teaching by the course instructor

| Anxiety_T | Anxiety impact on teaching (+/-/0)? | Anxiety_G |
| --- | --- | --- |

Writing quizzes, exams, or other graded assignments

| Anxiety_T | Anxiety impact on teaching (+/-/0)? | Anxiety_G |
| --- | --- | --- |

Not knowing the topic

| Anxiety_T | Anxiety impact on teaching (+/-/0)? | Anxiety_G |
| --- | --- | --- |

Answering student questions by email

| Anxiety_T | Anxiety impact on teaching (+/-/0)? | Anxiety_G |
| --- | --- | --- |

Time to prepare for teaching

| Anxiety_T | Anxiety impact on teaching (+/-/0)? | Anxiety_G |
| --- | --- | --- |

Time to complete a class activity

| Anxiety_T | Anxiety impact on teaching (+/-/0)? | Anxiety_G |
| --- | --- | --- |

Not knowing best teaching practices to use

| Anxiety_T | Anxiety impact on teaching (+/-/0)? | Anxiety_G |
| --- | --- | --- |

Grading

| Anxiety_T | Anxiety impact on teaching (+/-/0)? | Anxiety_G |
| --- | --- | --- |

Student behavior

| Anxiety_T | Anxiety impact on teaching (+/-/0)? | Anxiety_G |
| --- | --- | --- |

Teaching labs/discussion itself

| Anxiety_T | Anxiety impact on teaching (+/-/0)? | Anxiety_G |
| --- | --- | --- |

Advisor pushback/support

| Anxiety_T | Anxiety impact on teaching (+/-/0)? | Anxiety_G |
| --- | --- | --- |

| Anxiety_T | Anxiety impact on teaching (+/-/0)? | Anxiety_G |
| --- | --- | --- |

Seeing students outside of class

| Anxiety_T | Anxiety impact on teaching (+/-/0)? | Anxiety_G |
| --- | --- | --- |

| Anxiety_T | Anxiety impact on teaching (+/-/0)? | Anxiety_G |
| --- | --- | --- |

Meeting students for office hours

| Anxiety_T | Anxiety impact on teaching (+/-/0)? | Anxiety_G |
| --- | --- | --- |

Answering student questions in class

| Anxiety_T | Anxiety impact on teaching (+/-/0)? | Anxiety_G |
| --- | --- | --- |

Preparing to teach for an undergrad lab/class

| Anxiety_T | Anxiety impact on teaching (+/-/0)? | Anxiety_G |
| --- | --- | --- |

Being unable to answer a student’s question

| Anxiety_T | Anxiety impact on teaching (+/-/0)? | Anxiety_G |
| --- | --- | --- |

Speaking in front of the classroom

| Anxiety_T | Anxiety impact on teaching (+/-/0)? | Anxiety_G |
| --- | --- | --- |

Being evaluated by the course instructor/staff/faculty for your teaching

| Anxiety_T | Anxiety impact on teaching (+/-/0)? | Anxiety_G |
| --- | --- | --- |

**Cards for Research Anxiety (Yellow Cards)**

Meeting departmental deadlines

| Anxiety_R | Anxiety_G |
| --- | --- |

Leading my own project

| Anxiety_R | Anxiety_G |
| --- | --- |

Time management

| Anxiety_R | Anxiety_G |
| --- | --- |

| Anxiety_R | Anxiety_G |
| --- | --- |

Uncertainty of project success

| Anxiety_R | Anxiety_G |
| --- | --- |

| Anxiety_R | Anxiety_G |
| --- | --- |

Meeting your advisor

| Anxiety_R | Anxiety_G |
| --- | --- |

Leading lab meeting

| Anxiety_R | Anxiety_G |
| --- | --- |

Writing (e.g. proposals, grants, manuscripts)

| Anxiety_R | Anxiety_G |
| --- | --- |

Giving a seminar talk

| Anxiety_R | Anxiety_G |
| --- | --- |

Being reviewed by your advisor

| Anxiety_R | Anxiety_G |
| --- | --- |

Being unable to answer a question posed by a peer

| Anxiety_R | Anxiety_G |
| --- | --- |

Being unable to answer a question posed by my advisor/faculty

| Anxiety_R | Anxiety_G |
| --- | --- |

Living off your stipend

| Anxiety_R | Anxiety_G |
| --- | --- |

Attending lab meetings

| Anxiety_R | Anxiety_G |
| --- | --- |

Doing data analysis

| Anxiety_R | Anxiety_G |
| --- | --- |

Conducting lab/field work

| Anxiety_R | Anxiety_G |
| --- | --- |

Applying for grants, fellowships

| Anxiety_R | Anxiety_G |
| --- | --- |

Reading/understanding papers

| Anxiety_R | Anxiety_G |
| --- | --- |

Thinking about life after grad school

| Anxiety_R | Anxiety_G |
| --- | --- |

Interactions with lab mates

| Anxiety_R | Anxiety_G |
| --- | --- |

Being asked questions during a presentation

| Anxiety_R | Anxiety_G |
| --- | --- |

Interactions with faculty

| Anxiety_R | Anxiety_G |
| --- | --- |

| Anxiety_R | Anxiety_G |
| --- | --- |

**Supplemental Table 1:** Changes in each theme of anxiety related to **teaching** for each Biology GTA participant from 2016 to 2017. Anxiety themes identified for each participant in 2016 and 2017 were either kept (present each year), added (present only in 2017), resolved (present only in 2016), or were never present (not present in either year).

|  | **Impact on self** |  |  | **Impact on others** |  |  | **Lack of Self-efficacy** |  |  | **Role tension** |  |  | **Personal anxiety** | **I** |  |
| --- | --- | --- | --- | --- | --- | --- | --- | --- | --- | --- | --- | --- | --- | --- | --- |
| **Pseudonyms** | 2016 | 2017 | **Type of Change** | 2016 | 2017 | **Type of Change** | 2016 | 2017 | **Type of Change** | 2016 | 2017 | **Type of Change** | 2016 | 2017 | **Type of Change** |
| Hannah | **X** | **X** | Kept | **X** | **X** | Kept | **X** | **X** | Kept | **X** | **X** | Kept | **X** |  | Resolved |
| Cathy |  | **X** | Added | **X** | **X** | Kept | **X** | **X** | Kept |  | **X** | Added |  | **X** | Added |
| Raj | **X** |  | Resolved | **X** |  | Resolved | **X** |  | Resolved |  | **X** | Added |  |  | Never Had |
| Mark | **X** | **X** | Kept | **X** | **X** | Kept | **X** | **X** | Kept |  | **X** | Added | **X** | **X** | Kept |
| Emily | **X** |  | Resolved | **X** | **X** | Kept | **X** | **X** | Kept |  |  | Never Had | **X** |  | Resolved |
| Kayla | **X** |  | Resolved | **X** | **X** | Kept | **X** | **X** | Kept | **X** | **X** | Kept | **X** | **X** | Kept |
| Kaitlyn |  | **X** | Added | **X** |  | Resolved | **X** | **X** | Kept |  |  | Never Had |  | **X** | Added |
| Laretta | **X** | **X** | Kept | **X** | **X** | Kept | **X** | **X** | Kept |  |  | Never Had |  | **X** | Added |
| Raegan | **X** | **X** | Kept | **X** | **X** | Kept |  | **X** | Resolved | **X** | **X** | Kept | **X** | **X** | Kept |
| Madison | **X** | **X** | Kept |  | **X** | Added |  |  | Never Had |  | **X** | Added | **X** | **X** | Kept |
| William | **X** |  | Resolved | **X** |  | Resolved |  | **X** | Added |  | **X** | Added | **X** | **X** | Kept |
| Eric | **X** |  | Resolved |  | **X** | Added | **X** | **X** | Kept |  | **X** | Added | **X** | **X** | Kept |
| Lauren | **X** | **X** | Kept | **X** | **X** | Kept | **X** | **X** | Kept |  | **X** | Added |  |  | Never Had |
| Sarah | **X** |  | Resolved | **X** |  | Resolved | **X** |  | Resolved |  | **X** | Added | **X** |  | Resolved |
| Rebecca | **X** |  | Resolved | **X** | **X** | Kept |  | **X** | Added | **X** | **X** | Kept |  | **X** | Added |
| Samantha |  | **X** | Added |  | **X** | Added | **X** | **X** | Kept | **X** | **X** | Kept | **X** |  | Resolved |
| Julia | **X** | **X** | Kept | **X** | **X** | Kept |  | **X** | Added | **X** | **X** | Kept | **X** |  | Resolved |
| Arnold | **X** |  | Resolved | **X** | **X** | Kept | **X** | **X** | Kept | **X** | **X** | Kept |  |  | Never Had |
| Jose |  | **X** | Added | **X** |  | Resolved | **X** | **X** | Kept | **X** | **X** | Kept | **X** | **X** | Kept |
| Lucy | **X** | **X** | Kept | **X** |  | Resolved | **X** | **X** | Kept | **X** | **X** | Kept |  |  | Never Had |
| Anika | **X** |  | Resolved | **X** |  | Resolved | **X** | **X** | Kept |  |  | Never Had |  | **X** | Added |
| Jack | **X** |  | Resolved | **X** | **X** | Kept | **X** | **X** | Kept | **X** | **X** | Kept |  | **X** | Added |
| Sunny | **X** | **X** | Kept | **X** | **X** | Kept | **X** | **X** | Kept | **X** | **X** | Kept |  | **X** | Added |

**Supplemental Table 2:** Changes in each theme of anxiety related to **research** for each Biology GTA participant from 2016 to 2017. Anxiety themes identified for each participant in 2016 and 2017 were either kept (present each year), added (present only in 2017), resolved (present only in 2016), or were never present (not present in either year).

|  | **Impact on self** |  |  | **Impact on others** |  |  | **Lack of Self-efficacy** |  |  | **Role tension** |  |  | **Personal anxiety** |  |  |
| --- | --- | --- | --- | --- | --- | --- | --- | --- | --- | --- | --- | --- | --- | --- | --- |
| **Pseudonyms** | 2016 | 2017 | **Type of Change** | 2016 | 2017 | **Type of Change** | 2016 | 2017 | **Type of Change** | 2016 | 2017 | **Type of Change** | 2016 | 2017 | **Type of Change** |
| Hannah | **X** | **X** | Kept |  |  | Never Had | **X** | **X** | Kept | **X** | **X** | Kept | **X** |  | Resolved |
| Cathy | **X** | **X** | Kept |  |  | Never Had | **X** | **X** | Kept | **X** |  | Resolved |  |  | Never Had |
| Raj | **X** | **X** | Kept | **X** |  | Resolved | **X** | **X** | Kept |  |  | Never Had |  | **X** | Added |
| Mark | **X** | **X** | Kept | **X** | **X** | Kept | **X** | **X** | Kept |  |  | Never Had |  | **X** | Added |
| Emily | **X** | **X** | Kept | **X** | **X** | Kept | **X** | **X** | Kept |  |  | Never Had |  |  | Never Had |
| Kayla | **X** | **X** | Kept |  |  | Never Had | **X** | **X** | Kept |  |  | Never Had |  |  | Never Had |
| Kaitlyn | **X** | **X** | Kept | **X** |  | Resolved | **X** | **X** | Kept |  |  | Never Had |  |  | Never Had |
| Laretta | **X** |  | Resolved |  |  | Never Had | **X** | **X** | Kept |  |  | Never Had | **X** | **X** | Kept |
| Raegan | **X** | **X** | Kept | **X** |  | Resolved | **X** | **X** | Kept |  |  | Never Had | **X** |  | Resolved |
| Madison | **X** | **X** | Kept | **X** |  | Resolved | **X** | **X** | Kept |  |  | Never Had | **X** |  | Resolved |
| William | **X** |  | Resolved |  |  | Never Had | **X** | **X** | Kept |  |  | Never Had | **X** | **X** | Kept |
| Eric | **X** | **X** | Kept | **X** |  | Resolved | **X** | **X** | Kept |  | **X** | Added | **X** |  | Resolved |
| Lauren | **X** | **X** | Kept |  |  | Never Had | **X** | **X** | Kept |  |  | Never Had |  |  | Never Had |
| Sarah | **X** | **X** | Kept |  |  | Never Had | **X** | **X** | Kept |  | **X** | Added |  | **X** | Added |
| Rebecca | **X** | **X** | Kept |  | **X** | Added | **X** | **X** | Kept |  |  | Never Had | **X** |  | Resolved |
| Samantha |  | **X** | Added | **X** |  | Resolved | **X** | **X** | Kept |  |  | Never Had |  |  | Never Had |
| Julia | **X** | **X** | Kept |  | **X** | Added | **X** | **X** | Kept | **X** | **X** | Kept |  |  | Never Had |
| Arnold | **X** |  | Resolved | **X** |  | Resolved | **X** | **X** | Kept |  |  | Never Had |  |  | Never Had |
| Jose | **X** | **X** | Kept |  |  | Never Had | **X** | **X** | Kept | **X** | **X** | Kept | **X** |  | Resolved |
| Lucy | **X** |  | Resolved |  |  | Never Had | **X** | **X** | Kept | **X** | **X** | Kept |  |  | Never Had |
| Anika | **X** |  | Resolved | **X** |  | Resolved | **X** | **X** | Kept |  |  | Never Had | **X** |  | Resolved |
| Jack |  |  | Never Had |  | **X** | Added | **X** | **X** | Kept | **X** | **X** | Kept |  | **X** | Added |
| Sunny | **X** | **X** | Kept | **X** |  | Resolved | **X** | **X** | Kept | **X** | **X** | Kept |  |  | Never Had |

**Supplemental Table 3:** Demographics of Biology GTA interview participants as of the initial Fall 2016 interviews.

| **Pseudonyms** | **Gender** | **Ethnicity** | **Citizenship** | **Program** | **Teaching Experience Level** | **Career Aspirations** |
| --- | --- | --- | --- | --- | --- | --- |
| Hannah | Female | White | Domestic | PhD | Experienced | Non-Academic |
| Cathy | Female | White | Domestic | PhD | Experienced | Academic |
| Raj | Male | Non-White | International | PhD | Experienced | Academic |
| Mark | Male | White | Domestic | Master's | Experienced | Non-Academic |
| Emily | Female | White | Domestic | Master's | Novice | Academic |
| Kayla | Female | Non-White | International | Master's | Experienced | Academic |
| Kaitlyn | Female | White | Domestic | PhD | Novice | Non-Academic |
| Laretta | Female | White | Domestic | PhD | Experienced | Academic |
| Raegan | Female | White | Domestic | PhD | Novice | Academic |
| Madison | Female | White | Domestic | PhD | Experienced | Academic |
| William | Male | White | Domestic | PhD | Experienced | Non-Academic |
| Eric | Male | White | Domestic | PhD | Experienced | Non-Academic |
| Lauren | Female | White | Domestic | Master's | Novice | Non-Academic |
| Sarah | Female | White | Domestic | PhD | Novice | Non-Academic |
| Rebecca | Female | White | Domestic | PhD | Experienced | Non-Academic |
| Samantha | Female | White | Domestic | PhD | Experienced | Non-Academic |
| Julia | Female | White | Domestic | PhD | Novice | Non-Academic |
| Arnold | Male | White | International | PhD | Experienced | Academic |
| Jose | Male | Non-White | International | PhD | Novice | Academic |
| Lucy | Female | White | Domestic | PhD | Experienced | Academic |
| Anika | Female | Non-White | International | PhD | Novice | Non-Academic |
| Jack | Male | White | Domestic | PhD | Novice | Academic |
| Sunny | Female | Non-White | International | PhD | Novice | Non-Academic |
